# Prevalent Gut Phages Encode Modular Adhesins Mediating Epithelial Binding and Endoplasmic Reticulum Trafficking

**DOI:** 10.64898/2026.02.02.702849

**Authors:** Gábor Apjok, Tóbiás Sári, Orsolya Méhi, András Asbóth, Lilla Barna, Dóra Sala, Ilona Gróf, Bálint Márk Vásárhelyi, Szilvia Juhász, Csaba Pál, Péter Horváth, Ede Migh, György Schneider, Colin Hill, Mária Deli, Andrey Shkoporov, Bálint Kintses

## Abstract

Bacteriophages are crucial components of the human microbiome and hold promise as therapeutic agents. Yet, their physical interactions with mammalian cells remain poorly understood. Here, we developed a high-throughput platform to identify phages that adhere to epithelial layers and the proteins that mediate this interaction. The identified phages encode immunoglobulin (Ig)-like domain-containing proteins that, when displayed on a non-adherent phage, confer epithelial binding and internalization *in vitro,* and increased phage retention in the mouse gut *in vivo*. Phages encoding these adhesins are among the most abundant and prevalent human gut phages, including crAss-like phages and a distinct cluster of uncharacterized myophages. Domain sequence variation alters epithelial interaction profiles, and internalized phages traffic to the endoplasmic reticulum through the Golgi apparatus, suggesting access to non-degradative internalization pathways. These findings reveal widespread phage-human interactions in the human virome, with potential impacts on health and implications for next-generation phage therapeutics.

## Introduction

The mammalian microbiome is one of the most densely populated and complex ecosystems, comprising bacteria, archaea, viruses, and eukaryotes that collectively shape host physiology^1,2^. While bacteria are recognized as central to human health, bacteriophages, viruses that infect bacteria, are emerging as influential yet mechanistically underexplored components of host-associated microbial communities^1,3^. Despite their evolutionary distance, phages and human cells engage in complex interactions,^4,5^ with phage DNA and particles detected even in tissues once considered sterile, including blood, kidney, cerebrospinal fluid, and the brain^5-8^. Phages can persist in the body for extended periods^5-8^, synergize with the immune system^9-11^, and even enhance human cell viability among other effects^12-15^. However, the molecular determinants that enable specific phages to engage with mammalian cell surfaces and cellular pathways remain largely unknown.

Epithelial layers form the primary barrier between the human body and its environment. To interact with it, microbes have evolved diverse strategies^16-21^, particularly in the gastrointestinal tract, where complex tripartite interactions unfold between bacteria, phages, and host epithelial cell surfaces^1^. It is estimated that as many as 31 billion phage particles may cross the gut barrier daily^22^, yet the mechanisms and consequences of this translocation are unclear. Several studies suggest that phages displaying surface-exposed adhesion modules can bind mucosal components, thereby increasing residence time at epithelial interfaces^23-25^. Among these, immunoglobulin (Ig)-like domains have been proposed as key determinants, and early genomic surveys revealed their widespread occurrence in phage genomes^5,26^. However, the rapid expansion of available phage genome sequences necessitates a systematic reassessment^27^. Critically, most Ig-like domains and other potential adhesins remain functionally uncharacterized, making it unclear which phages can actively adhere to epithelial surfaces, which domains are responsible, and whether such interactions extend beyond mucus binding to include direct engagement with epithelial cells. A notable exception is the T4 phage, whose capsid surface protein, <u>H</u>ighly immunogenic <u>o</u>uter <u>c</u>apsid (Hoc), binds to intestinal mucus, forming the basis of the <u>B</u>acteriophage <u>A</u>dherence to <u>M</u>ucus (BAM) hypothesis^28^. As a result of adhesion, the eukaryotic cells can internalize the T4 phage *in vitro*^29-31^. Whether similar mechanisms operate in dominant but largely uncultivated gut phages, such as crAss-like phages and other uncharacterized phage groups, remains unknown^27^. Systematically uncovering these mechanisms is crucial for understanding the physiological effects of phages on the human host and for advancing the development of novel phage therapeutics.

Here, we developed a high-throughput pipeline to systematically identify phages that adhere to epithelial layers and the phage-encoded proteins mediating these interactions. Specifically, by phage affinity panning on epithelial surfaces, virome sequencing, domain-centric analyses, and functional validation through phage engineering, we demonstrate that modular Ig-like domain-containing proteins confer epithelial binding and uptake *in vitro* when displayed on a non-adherent phage scaffold. Importantly, the displayed proteins also increase phage retention in the murine gastrointestinal tract *in vivo*, linking epithelial association to altered phage persistence under physiological conditions. We further show that phages encoding the identified Ig-like domains are highly abundant and prevalent in the human gut virome and that such phages carry the adhesins in different combinations, indicating that they precisely control adherence strength and binding partners on the mucosal surfaces. Finally, fluorescence microscopy reveals that internalized phages can traffic to the endoplasmic reticulum, suggesting access to non-degradative intracellular pathways. These findings reveal widespread phage-human interactions with implications for human health and targeted phage delivery.

## Results

### High-Throughput Identification of Adhering Phages and their Adhesins

To capture diverse phages for the assay, we pooled fecal viral filtrates from ten healthy and ten dysbiotic individuals (Fig. 1A), leveraging substantial inter-individual and health-associated variability in the phageome^32-34^. We spiked the filtrate with the well-characterized T4 phage as a positive control due to its known mucosal adhesion and intracellular entry via macropinocytosis ^13,28^. Batches of the filtrate were applied to either low mucus-producing Caco-2 or high mucus-producing HT29-MTX cell layers to assess phage adhesion to the glycocalyx, the overlaying mucus, or other epithelial cell surface receptors^35^ (Fig. 1A). Following incubation, non-associated phages were removed by repeated, gentle rinsing of the sample and collected as the supernatant (“effluent” fraction). Adhering phages were subsequently eluted from the cell layer (“residual” fraction) with an established methodology^7^ (see Methods). Phage DNA was extracted from both fractions and the initial filtrate, and subjected to next-generation sequencing. The entire procedure was performed in two biological replicates to support reproducibility (see Methods).

**Figure 1.**
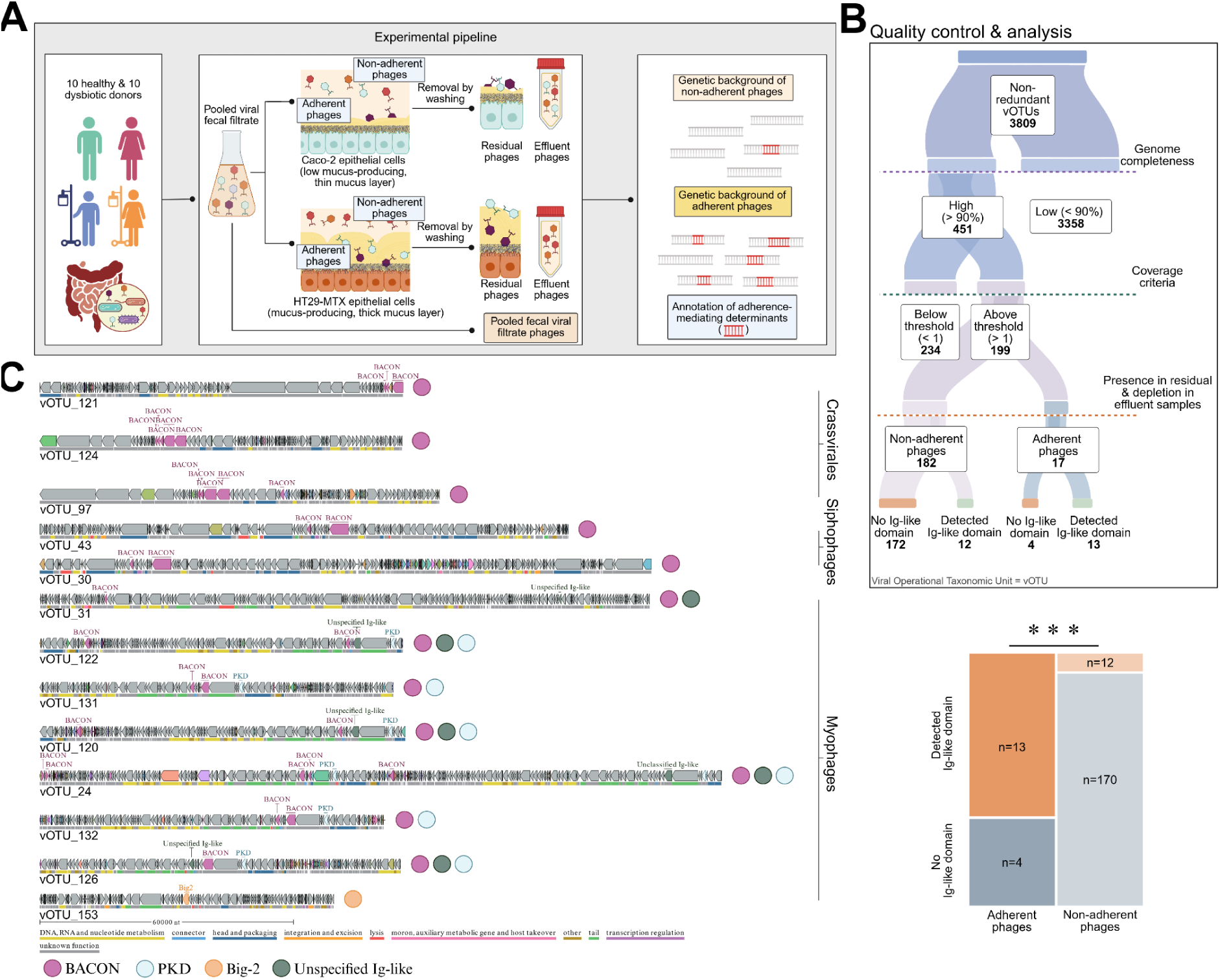
High-throughput identification of adherent phages and their proteins involved in gut epithelial adherence and uptake. **A)** Overview of the experimental workflow used to separate phages that adhere to epithelial cells (residual fraction) from non-adherent phages (effluent fraction), followed by short-read sequencing and downstream analyses. **B)** Quality control and analytical steps used to identify adherent and non-adherent viral operational taxonomic units (vOTUs) and their adhesins, including quality filters (genome completeness and coverage), adherence phenotype (see methods), and the enrichment of adhesins. Bold numbers indicate the number of vOTUs retained at each step. Ig-like domain-containing ORFs are significantly enriched in adherent compared with non-adherent vOTUs (two-sided Fisher’s exact test, p = 0.00001, N=199). **C)** Schematic representation of predicted open reading frames (ORFs) in 13 adherent vOTUs encoding Ig-like domains. Ig-like domains are labeled by domain type, and annotated phage proteins are color-coded at the bottom of each genome. All data are available in Tables S1-S5.

Assembly of sequencing reads yielded 97,460 contigs of phage origin, clustered into 3,809 viral Operational Taxonomic Units (vOTUs) at 95% nucleotide identity (Table S1). To ensure accurate detection of complete domain repertoires, vOTUs were filtered for quality based on genome completeness (≥90%) and coverage (≥1), resulting in a curated set of 199 high-confidence vOTUs (Fig. 1B, Methods). Of these, 17 were classified as adherent and 182 as non-adherent based on two stringent criteria: (i) presence in all four residual samples (two biological replicates across both epithelial cell types), indicating adherence and (ii) concurrent depletion from all four effluent samples, indicating cellular uptake (Fig. 1B, Methods; Table S2). The remaining 182 vOTUs were classified as non-adherent. Finally, to identify candidate protein mediators of phage adherence and uptake, we performed a genome-wide survey of phage-encoded protein domains in the 199 complete vOTUs. Specifically, protein domains were systematically evaluated for enrichment in adherent versus non-adherent vOTUs and then screened for putative adherence-related functions based on prior literature (Table S3, S4; Methods).

In agreement with the bacteriophage adherence to mucus (BAM) model^28,36^, adherent vOTUs encoded Ig-like-domain-containing open reading frames (ORFs) disproportionally more frequently than non-adherent vOTUs (p < 0.00001, Fisher’s exact test, Fig. 1B, Table S2). Interestingly, the thirteen Ig-like-bearing adherent vOTUs contained multiple such domains, totaling 45 ORFs encoding 67 Ig-like regions (Table S5). Of these ORFs, 79% were located proximal to structural proteins, suggesting their incorporation into structural elements (Fig. 1C, Table S5). Classification of the 67 Ig-like regions revealed four major Ig-like domain types, with the **B**acteroidetes **A**ssociated **C**arbohydrate-binding **O**ften **N**-terminal (BACON) domain representing approximately 64% of all identified Ig-like regions (Fig. 1C, Table S5). They are present in twelve of the thirteen Ig-like domain-encoding adherent vOTUs (Fig. 1C, Table S5, S6), despite originating from multiple, evolutionarily distinct phage groups, as revealed by the network analysis of shared protein orthogroups (Fig. S1, Table S7). The two largest BACON-encoding phage clusters were crAss-like phages and an unspecified phage group with myovirus morphology (Fig. S1, Table S1 and S7). Beyond the unspecified Ig-like domains, the second most prevalent type was the polycystic kidney disease (PKD) domain, present in T4 Hoc protein and in polycystin-1 cell-surface glycoproteins, hypothesized to mediate protein-protein and protein-carbohydrate interactions^37^. PKD domains occurred in ∼46% (6/13) of Ig-like-domain bearing adherent vOTUs, including all vOTUs from the unspecified myophage group (Fig 1C; Fig. S1, Table S6). Based on CRISPR spacer match predictions, these vOTUs infect *Bacteroidota* hosts, akin to crAss-like phages (Tables S1, S2 and S8). As an exception to the general trend of encoding multiple Ig-like domains, one vOTU (vOTU_153) contained only a single Ig-like domain, classified as a Big2-type Ig-like domain (Fig 1C, Fig. S1, Table S6).

### Displaying transferred adhesins facilitates phage-epithelial interaction

To validate the predicted function of the identified Ig-like domain-containing proteins, we displayed a subset of them on the surface of the non-adherent *Escherichia coli* phage K1F scaffold. Three candidate ORFs PCR-amplified from the fecal viral filtrate were integrated into the K1F genome to create an occasional translational fusion with the capsid protein gp29 (Fig. S2, Table S9, and Methods). This design ensures that the fusion protein is displayed only on a fraction of the capsid-forming proteins, minimizing disruption to phage assembly and function^38^. This resulted in three engineered phages: K1F::S, carrying a protein with a single BACON domain, detected in vOTU_120, 122, and 131 (all three of which are uncharacterized myovirus phages, Fig. 1C); K1F::L, encoding a protein with one BACON and one unspecified Ig-like domain, detected in vOTU_131; and K1F::7649, encoding a closely related variant of the previous protein, differing by only two amino acids, detected also in vOTU_131 (Table S9). As a positive control, we generated a K1F phage variant displaying the mucus-binding Hoc protein of T4^36^, designated as K1F::hoc, encoding two PKD and one unspecified Ig-like domains (Table S9). Together, these four proteins cover the three most prevalent, potentially adherence-mediating domain types (BACON, PKD, and unspecified Ig-like domains) identified in adherent vOTUs (Table S6). Characterization of infection dynamics confirmed that the viability of K1F remained unaffected by the displayed Ig-like domains (Fig. S3, Table S9). Wild-type K1F, engineered K1F variants, and wild-type T4 phage were fluorescently labeled using SYBR™ Gold and applied to epithelial cell monolayers for quantitative association and uptake assays (Methods). Three human epithelial cell lines with distinct surface properties were examined: Caco-2 and HT29-MTX intestinal epithelial cells, which display low basal uptake potential, and A549 lung epithelial cells, which exhibit high mucus production and elevated endocytic activity^29,39,40^. Intracellular fluorescent signal changes were tracked over 18 hours in three replicates (see Methods)

While wild-type K1F showed minimal internalization in the tested epithelial cell types, all four recombinant phages demonstrated significantly higher uptake with varying rates (Fig. 2ABC, Table S11). High mucus-producing cell lines (A549 and HT29-MTX) exhibited higher internalization of K1F displaying Ig-like domains compared to the low mucus-producing Caco-2 line, regardless of differences in non-receptor-mediated endocytic activity of A549 and HT29-MTX (Fig. 2D). The magnitude and kinetics of signal accumulation varied among constructs, indicating that distinct Ig-like domain architectures confer different interaction strengths. The T4 phage showed a significantly higher internalization rate than the engineered K1F::hoc phage across all three cell types (Fig 2C). This discrepancy may be explained by (i) the native T4 phage carrying approximately three times more Hoc proteins than the engineered K1F variant,^41-43^ (ii) the C-terminal capsid binding domain of the Hoc protein theorized to specifically interact with the native T4 capsid proteins and its transfer to the K1F phage perturbed its native conformation^44^ or (iii) a Fibronectin type III Ig-like domain on the T4 phage tail fiber may further facilitate interaction with eukaryotic cell surface residues beyond the Ig-like domains of the Hoc protein (Table S3).

**Figure 2.**
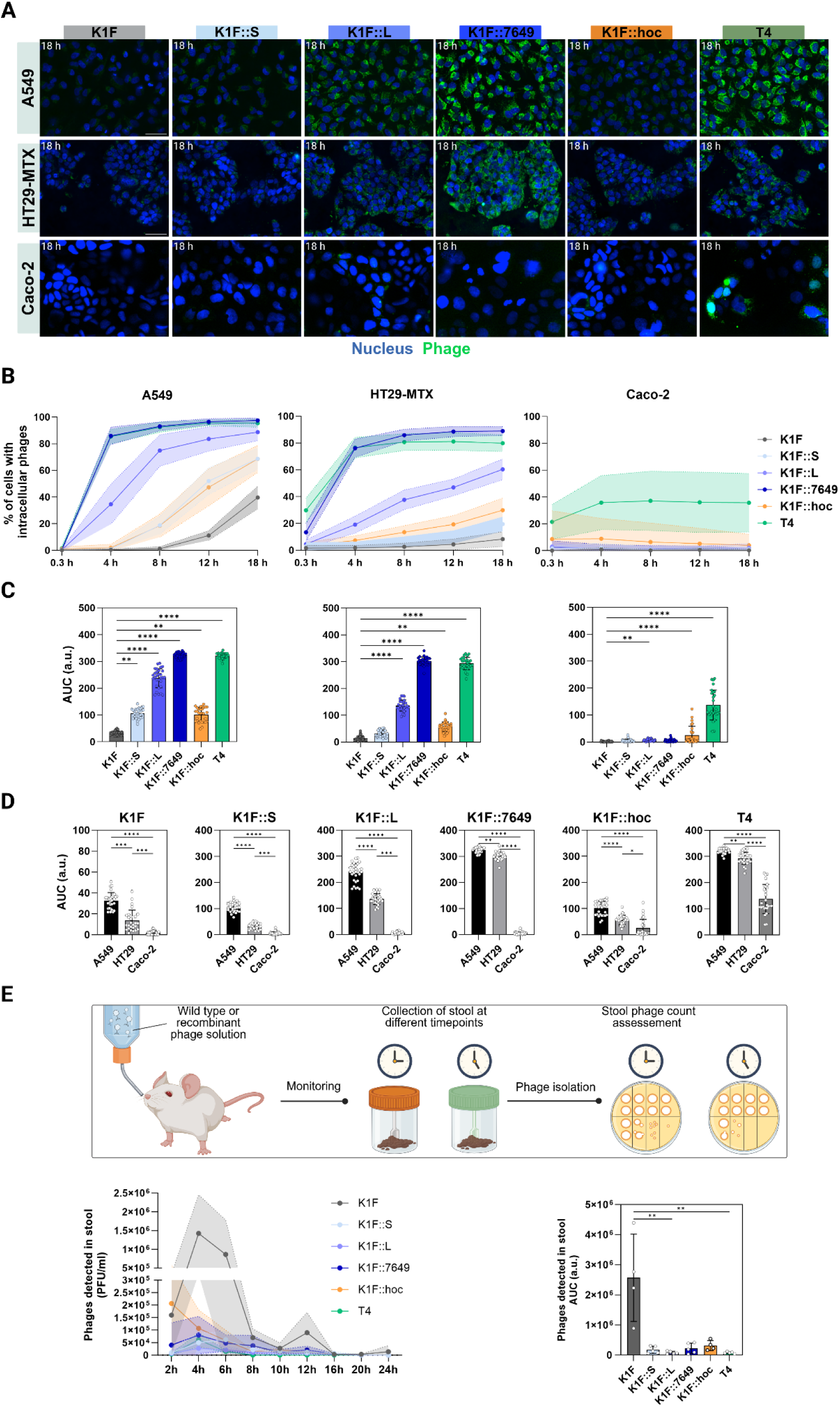
Adhesin display enhances epithelial adherence of K1F phage *in vitro* and *in vivo*. **A)** Confocal (A549, HT29-MTX) and widefield (Caco-2) microscopy images showing internalization of wild-type and adhesin-displaying K1F phages, as well as wild-type T4 phage, in A549, HT29-MTX, and Caco-2 cell lines after 18 h of co-incubation. Nuclei were stained with Hoechst (blue) and phages with SYBR™ Gold (green). Z-stacks were acquired across the full nuclear depth from apical to basal planes (see Methods). Scale bar: 50 µm. **B)** Time-course quantification of cellular uptake of wild-type and adhesin-displaying K1F phages and T4 phage in A549, HT29-MTX, and Caco-2 cells over 18 h (N = 27). The percentage of cells containing intracellular phages was calculated from 27 fields of view. Mean ± SD are shown. **C)** Area under the curve (AUC) values calculated from the uptake curves in B. Each dot represents one field of view (N = 27); mean ± SD are shown. P values were calculated using the Kruskal-Wallis test (two-tailed). a.u., arbitrary units. **D)** Comparison of phage uptake by high mucus-producing A549 and HT29-MTX cells and low mucus-producing Caco-2 cells, quantified as AUC values derived from B. Each dot represents one field of view (N = 27); mean ± SD are shown. Significance was assessed by Kruskal-Wallis test (two-tailed)(*p < 0.05, **p < 0.01, ***p < 0.001, ****p < 0.001). **E)** Time-course and AUC analysis of phage counts detected in mouse stool, measured as shown in the schematic. Each dot represents one animal (N = 5, except K1F and K1F::hoc, where one animal per group was excluded).Significance was assessed by Kruskal-Wallis test (**p < 0.01). Data are available in Table S11 for B, C and D and Table S12 for E.

### Adhesins increase retention in the murine gastrointestinal tract

To assess whether the identified phage-host interactions also occur under physiological conditions, we examined whether displaying Ig-like domains on the K1F phage affects phage retention in the mouse gut. To this end, mice were gavaged with 2 x10^8^ PFU of phages, and fecal shedding was monitored over 24 hours (Methods). Wild-type K1F and T4 phages were used as negative and positive controls for adherence, respectively.

Consistent with previous reports^45^, wild-type K1F phage exhibited rapid gastrointestinal transit with fecal phage counts peaking at approximately 4 hours post-administration (Fig. 2E, Table S12). In contrast, engineered K1F variants displaying Ig-like domain-containing proteins showed reduced fecal recovery over time relative to wild-type K1F, indicating altered gastrointestinal retention dynamics. Among these, K1F::L displayed the most pronounced reduction in fecal shedding, comparable to that observed for T4 phage. The remaining engineered variants exhibited intermediate phenotypes, mirroring the trends observed i*n vitro*. Overall, displaying the Ig-like containing proteins reduced the clearance of the recombinant phages from the animals. This observation suggests sequestration into the mucin layer with possibly internalization by epithelial cells, thereby altering their elimination from the body.

### Phages with modular adhesins are abundant and prevalent in the healthy human virome

Next, we asked whether phages encoding the experimentally identified Ig-like domains represent rare curiosities or adherence to mucosal surfaces is a broadly applied phenomenon in the human virome. We analysed two datasets: (i) Cenote Human Virome Database^46^, encompassing seven human body sites from nearly 6,000 metagenomes, comprising >45,000 vOTUs, designated here as the Broad Human Virome; and (ii) faecal viromes of 40 healthy individuals and 39 patients with Crohn’s disease or ulcerative colitis, consisting of >28,000 vOTUs, designated here as 79-Individual Gut Virome^47^.

Across the Broad Human Virome and the 79-Individual Gut Virome datasets, 10.1% and 5.5% of vOTUs, respectively, harboured at least one of the four Ig-like domain types that we identified in the adherent vOTUs (BACON, PKD, unspecified Ig-like, Big-2), Table S13, S14). Among these, we identified 162 vOTUs encoding 475 Ig-like domain-containing proteins that share ≥80% sequence identity with the 45 ORFs detected in the 13 experimentally identified adherent vOTUs (Fig. 3A, Supplementary file 1, Table S15, S16). Given the high sequence similarity, these proteins are also likely mediators of adherence. The majority of them are modular, encoding BACON, PKD, and unspecified Ig-like domains in different combinations (Fig. 3A). CRISPR spacer matches suggest that the vOTUs encoding them infect hosts belonging to the *Bacteroidota* phylum (Table S13), which are commonly associated with mucosal niches and utilization of host-derived glycans^48^. This implies that these phages attain persistence and co-localization with their hosts by adhering to mucus.

**Figure 3.**
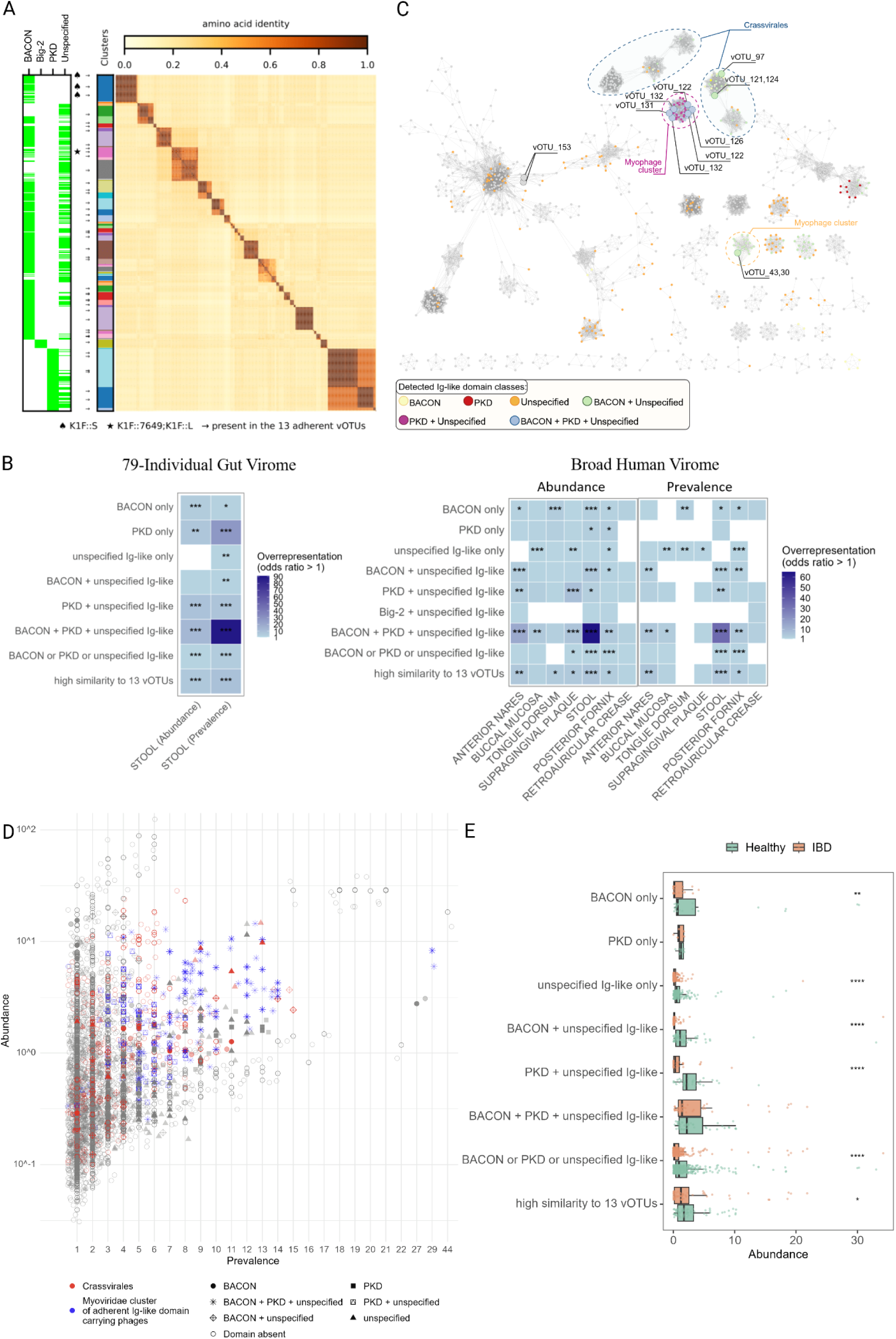
Genomic distribution of Ig-like domains in the human virome. **A)** Clustering of 475 Ig-like domain-containing ORFs from the Broad Human Virome and the 79-Individual Gut Virome that share ≥80% sequence identity with the 45 Ig-like domain-containing ORFs identified in 13 experimentally validated adherent vOTUs. ORFs were clustered at ≥80% identity and coverage. Arrows indicate ORFs with 100% sequence identity to those in the experimentally identified vOTUs. In the left panel, green lines denote the presence of Ig-like domain types. Symbols indicate ORFs present in K1F::S, K1F::L, and K1F::7649, respectively. **B)** vContact3 protein-sharing network of high-quality phage vOTUs (N = 2269) from the 79-Individual Gut Virome. Nodes and edges represent vOTUs and shared protein clusters, respectively. Colored and grey nodes indicate vOTUs with and without Ig-like domains, with domain types color-coded. Dashed outlines highlight major clusters enriched for Ig-like domain-encoding phages, including crAssvirales and a distinct cluster of uncharacterized myophages. Selected adherent vOTUs from the epithelial adherence screen are labeled. **C)** Phages harboring BACON, PKD, or unspecified Ig-like domains are overrepresented among highly abundant and prevalent phages across both datasets. Blue intensity indicates the degree of overrepresentation (odds ratio >1) based on two-tailed Fisher’s exact test. N is provided in Tables S17 and S18, p-values were Benjamini-Hochberg corrected. **D)** Abundance versus prevalence of vOTUs in the 79-Individual Gut Virome. vOTUs of interest are distinguished by shape. Uncharacterized myoviruses and CrAss-like phages are shown in red and blue, respectively (N = 2036 from 79 individuals). **E)** Abundance of vOTUs encoding different Ig-like domains in healthy individuals versus IBD patients in the 79-Individual Gut Virome (two-tailed Mann-Whitney U test, N = 1544 from 40 healthy and 39 IBD individuals, Table S14). Significance levels: *p < 0.05, **p < 0.01, ***p < 0.001, ****p < 0.0001.

We next tested a key prediction of the BAM model, namely, whether Ig-like domain-encoding phages are more abundant and prevalent in mucosal body sites compared with phages that do not encode these domains^28^. To ensure unbiased comparisons, we restricted our analysis to high-quality phage genomes (>90% completeness and high or medium confidence). Among these genomes, BACON, PKD, or unspecified Ig-like domains were present in 12.8% and 15.6% of the vOTUs in the Broad Human Virome and the 79-Individual Gut Virome, respectively (Table S13, S14). Most importantly, these phages were strongly enriched among abundant and prevalent vOTUs across both datasets, with the exception of the only body site not covered by a mucosal surface, the retroauricular crease (the skin area behind the ear, Fig. 3B; Tables S17, S18; Methods). The enrichments were most pronounced for phages encoding BACON, PKD, and unspecified Ig-like domains together (Fig. 3C). Importantly, when we controlled for host-driven ecological variations by analysing only Bacteroidota-infecting vOTUs, the enrichments of these Ig-like domain-carrying vOTUs in the high abundance and prevalence categories remained similarly strong (Fig. S4, Table S19). Generating a protein-sharing network from the 79-Individual Gut Virome revealed that vOTUs encoding these domain types cluster across the phage network (Fig. 3C), consistent with the clustering of the experimentally identified adherent vOTUs (Fig. S1). These phage groups were, again, *Crassvirales* and uncharacterized clusters of myophages (Fig. 3C). Notably, individual vOTUs from the largest uncharacterized myovirus group that harbor all three domain types were detected, on average, in 8 out of the 79 individuals, placing 82% of these 50 vOTUs among the top 10% most prevalent human gut phages (Fig. 3D). All 50 vOTUs showed >90% nucleotide identity with reference members of the Flandersviridae, strongly indicating that they belong to this phage family. CrAss-like phages utilizing these domains are similarly abundant and prevalent in the 79-Individual Gut Virome (Fig. 3D). These findings indicate that by promoting prolonged retention in the gut environment, mucin binding indirectly raises the likelihood that these phages appear consistently across individual phageomes.

Finally, we examined whether vOTUs carrying at least one BACON, PKD, or unclassified Ig-like domain were associated with healthy or disease-associated microbiomes (Methods). This analysis leveraged association data between phages from the Broad Human Virome and 11 different diseases^46^. Adherent vOTUs were not enriched in any disease category (Table S20) but were instead more abundant in the gut virome of healthy individuals than in those of patients with inflammatory bowel disease (IBD) (Fig. 3E), consistent with their dependence on an intact mucosal architecture.

### Minimal Ig-like Domain Variations Alter Epithelial Surface Targeting

To dissect how Ig-like domain sequence variation influences epithelial interaction partners, we examined whether the identified adhesins differ in their sensitivity to perturbations of epithelial surface components. We focused on high mucus-producing HT29-MTX cells that were subjected to treatments targeting distinct cell surface structures. Specifically, epithelial cultures were treated with N-acetylcysteine (NAC) to disrupt disulfide bond-mediated mucin structure^49^, cyclosporin A (CSA) to induce mucin aggregation^49-51^, or heparinase I to enzymatically remove heparan sulfate moieties from both the mucus layer and the underlying glycocalyx^52-54^. The impact of these treatments on cellular phage uptake was evaluated by investigating fluorescently labeled phage internalization on treated and untreated HT29-MTX cell cultures.

All cell surface treatments significantly reduced internalization of all the engineered K1F phages (Fig. 4, Table S21). However, while uptake of K1F::S, K1F::7649, and K1F::hoc was nearly abolished by all treatments, the disruption of the mucus layer with NAC and CSA had a notably smaller effect on K1F::L. Cleaving heparan sulfate further diminished K1F::L binding (Fig. 4 and Fig. S5, Table S21), suggesting that K1F::L uniquely interacts with both mucin and membrane-bound proteoglycans. This distinction between K1F::L and K1F::7649 is striking, as the two Ig-like domains differ by only two amino acids (Table S9).

**Figure 4.**
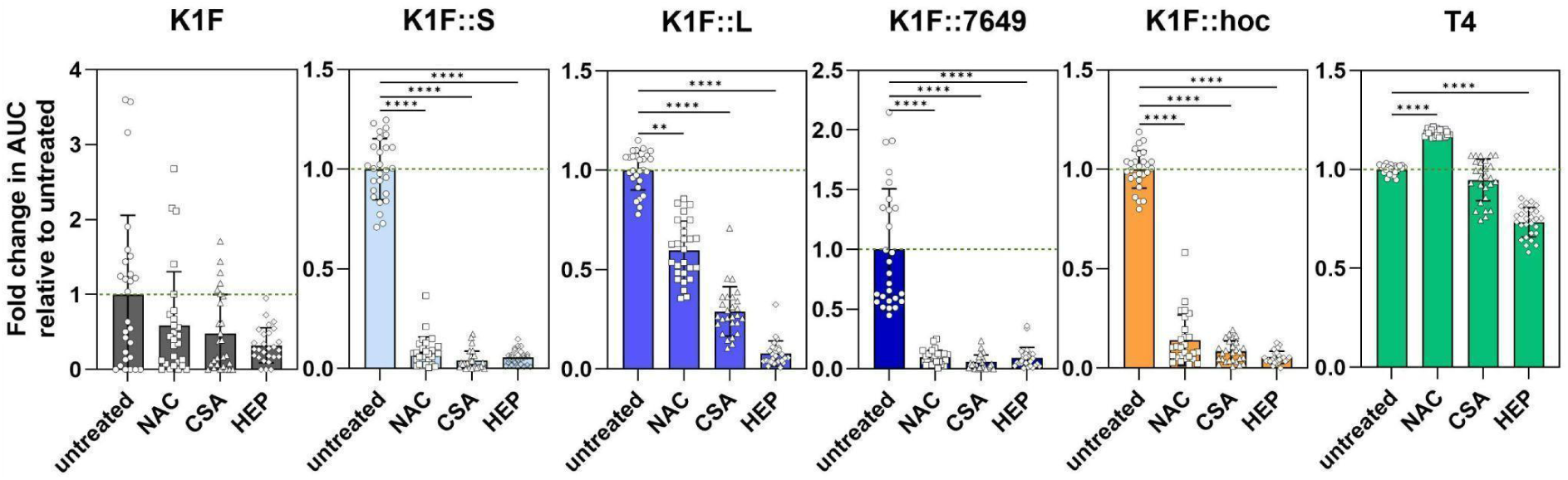
Effects of cell surface disruptions on the internalization of Ig-like domain-displaying K1F phages. Plots show the internalization of wild-type and engineered K1F phages, as well as T4 phages into the HT29-MTX cell line following different cell surface treatments. Internalization is expressed as the relative fold change in area under the curve (AUC), normalized to the untreated control (set to 1, indicated by a green dotted line). AUC was calculated from the percentage of cells with intracellular phages. Mean and standard deviation are shown. Each dot represents one field of view (N = 27). Stars indicate p-values calculated from Kruskal-Wallis test (two-tailed)(** p < 0.01, **** p < 0.001). Abbreviations: NAC - N-acetylcysteine, CSA - cyclosporin A, HEP - heparinase I. Data is available in Table S21)

Overall, adhering phages may bind to mucin and to the glycocalyx depending on the sequence of the Ig-like domain. Specific Ig-like domains that bind directly to the epithelial surface may enable internalization even in mucin-depleted or ulcerated sites where the cell surface is exposed.

### Phages with Adhesins Accumulate in ER, Evading Degradative Pathways

Phage internalization by eukaryotic cells can occur through various routes - macropinocytosis-, phagocytosis-, receptor-mediated clathrin or caveolin endocytic pathways, and less conventional pathways - depending on cell type and phage-surface interactions (Fig. 5A)^5,23,55,56^. To investigate intracellular trafficking of the engineered phages, we determined their colocalization with different organelles following entry into A549 epithelial cells. Prior to phage exposure, cells were labeled with fluorescent markers specific for lysosomes, the Golgi apparatus, or the endoplasmic reticulum (ER), and confocal imaging was performed to quantify colocalization with fluorescently labeled phages over time (Methods).

**Figure 5.**
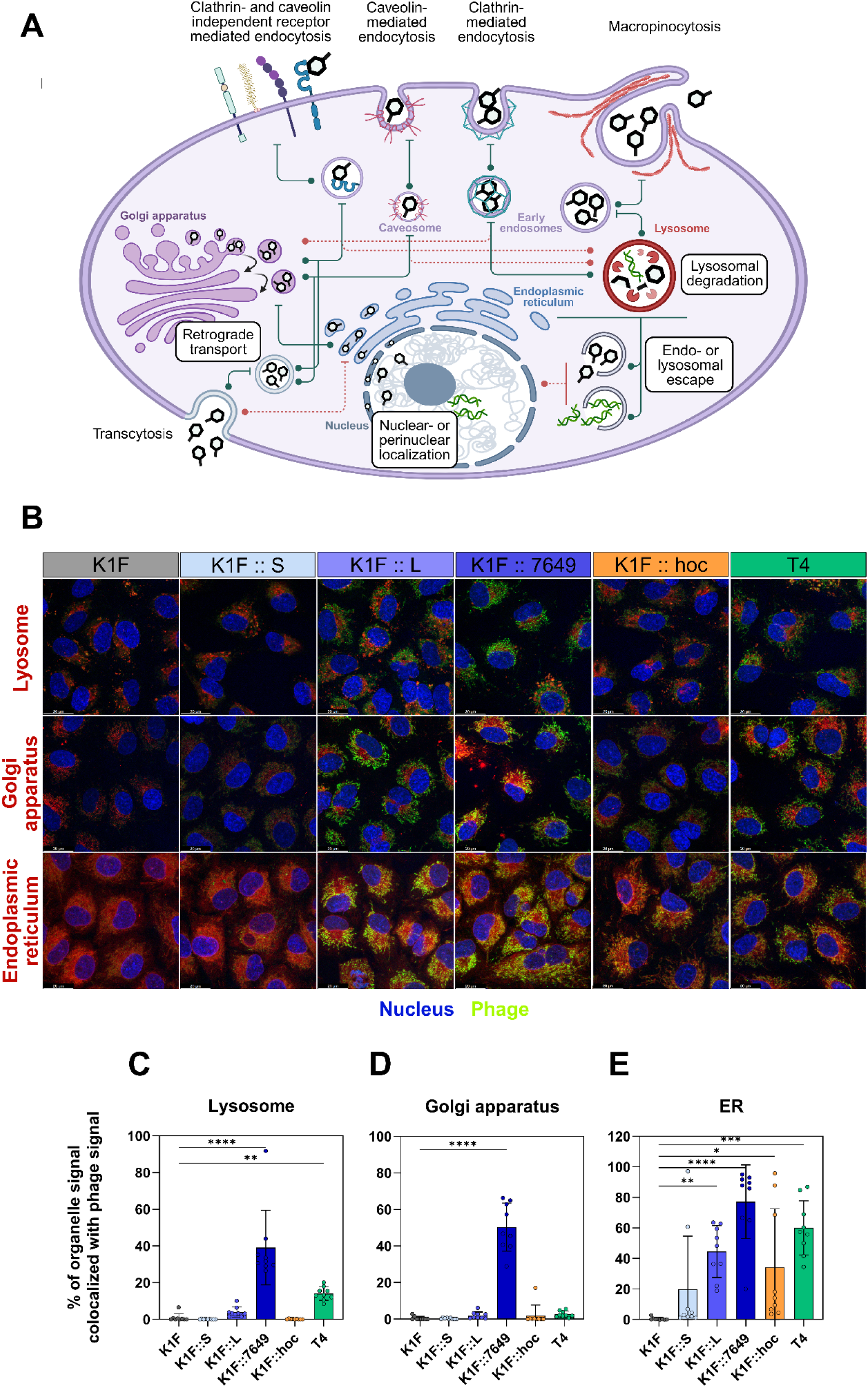
Potential internalization routes of phages and the colocalization patterns of engineered phages with subcellular compartments. **A)** Schematic illustration of potential entry and trafficking pathways for phages interacting with human cells, based on known endocytotic routes. Conventional routes are shown in green, while less common directions are marked by dashed red lines. **B)** Confocal microscopy images showing the colocalization of labeled wild-type and engineered phages (green: SYBR™ Gold dye) with lysosomes, the Golgi-apparatus, and the ER (red: Lysotracker^TM^ Deep Red, BODIPY^TM^ TR Ceramide, and ER-Tracker^TM^ Red dyes, respectively) Scale bar, 20µm. **C-E)** Colocalization of phages with different organelles: lysosome **(C)**, Golgi-apparatus **(D),** and ER **(E)** after six hours of incubation. Colocalization was quantified by the percentage of organelle-specific red signals overlapping with phage-specific green signals based on Manders’ overlapping coefficient. Mean and standard deviation are shown. Each dot represents one field of view. Stars indicate p-values calculated from Kruskal-Wallis test (two-tailed, N = 9, * p < 0.05, ** p < 0.01, *** p < 0.001, **** p < 0.001).

Engineered phages exhibited a distinct and reproducible intracellular localization pattern. Within 6 hours of exposure, all engineered variants consistently displayed strong colocalization with all three markers, implying multiple uptake- and intracellular traffic-routes available for the modified K1F phages (Fig. 5B-E, Fig. S6-8, Table S22). The level of accumulation within the cell organelles correlated with phages’ internalization efficiency. Specifically, K1F::L and K1F::7649, the two phages with the highest uptake, demonstrated the most colocalized phage with the Golgi and ER. These results support a model in which adhering phages enter epithelial cells through receptor-mediated internalization, partially evade canonical degradative processing, and traffic preferentially to the ER, likely via a rapid endosome-to-Golgi-to-ER pathway. The ER appears to act as a terminal destination under these experimental conditions.

In summary, engineered phages access conserved, non-lysosome-dominated intracellular routes with the ER as a prominent accumulation site, either by subverting host sorting processes or by engaging retrograde-competent endocytic pathways.

## Discussion

Our study established that physical interactions between bacteriophages and the human epithelium are widespread and an ecologically consequential feature of the gut virome. We demonstrate this by identifying phages that bind to epithelial cells and pinpoint to adhesins that mediate these interactions (Fig. 1). When three of these Ig-like domain-containing adhesin candidates, together with the PKD domain-containing Hoc protein from bacteriophage T4 as control, were displayed on the otherwise non-adherent *Escherichia coli* phage K1F, the engineered phages exhibited increased epithelial adherence and uptake *in vitro* across multiple epithelial cell types and increased gut retention in mice *in vivo* (Fig. 2). Phages encoding closely related adhesins are predominantly crAss-like phages and previously uncharacterized myophages which rank among the most abundant and prevalent phages in the human gut virome (Fig. 3). The likely attachment sites include the cell membrane-embedded glycocalyx layer and the overlying mucus (Fig. 4), consistent with the binding behavior of PKD-domain phages such as T4 phage^24,57,58^. Following epithelial uptake, the internalized phages localize mainly to the Golgi and the endoplasmic reticulum (ER), bypassing classical degradative compartments such as lysosomes (Fig. 5). Given that intracellular trafficking routes into the ER are typically non-degradative, it is plausible that internalized phages remain structurally intact within this compartment; however, this was not experimentally investigated. Such routing mirrors retrograde trafficking pathways exploited by certain bacterial toxins and non-enveloped viruses, raising the possibility that phage surface features bias intracellular sorting decisions following entry^59^.

These findings reveal that phage-epithelial interactions are far more widespread and complex than previously appreciated, with potential consequences for both the phage and the human host. Our data provide direct experimental support for a central premise of the bacteriophage adherence to mucus (BAM) model that adhesion to mucosal surfaces confers a selective advantage by spatial co-localization with bacterial hosts and increased phage retention which results in high abundance and prevalence in the gut environment^25,28,36^. Importantly, our results extend this concept beyond individual model phages to a diverse set of abundant and prevalent gut phages, including crAss-like phages and previously uncharacterized myoviruses. The independent emergence of similar modular adhesins within distinct evolutionary lineages further indicates that epithelial and mucus adhesion is a dominant and repeatedly selected ecological strategy in the human gut virome. The high prevalence of adherent phages in healthy gut viromes and the depletion in IBD suggests that adhesion-competent phages are signature components of a well-functioning mucosal environment. Their reliance on stable bacterial hosts and preserved mucus architecture explains their depletion when these conditions break down during inflammation, underscoring that their distribution reflects ecosystem health rather than a pathogenic role.

Phage adherence and internalization also hold promise for innovative therapeutic applications: engineered adhesins on the phage surfaces could enhance tissue targeting, retention, or cellular uptake, potentially enabling intracellular delivery or immune modulation. The observed ER trafficking raises the prospect of using phages to deliver therapeutic payloads to cells affected by ER-related disorders such as cystic fibrosis^60^, Fabry disease^61^, or Gaucher disease^62^. Nevertheless, understanding these adhesins highlights the importance of screening therapeutic phages for human-interaction domains to prevent unintended interactions with host tissues.

Several limitations of our study remain. In our high-throughput adherence assay, high amounts of remnant DNA from eukaryotic cells reduce assay sensitivity, making it likely that low-abundance adherent phages remain undetected. We do not yet know under which physiological or pathological conditions epithelium-associated phages undergo internalization in the human gut *in vivo*, where mucus architecture, workings of the immune system, and epithelial turnover differ substantially from in vitro systems. The functional integrity of internalized phages by the epithelial cells also remains unknown, as do the immunological consequences of their entry into host cells. Future studies should address these aspects and explore the long-term outcomes of these phage-human interactions. Nonetheless, our work identifies and validates modular genetic determinants of phage adherence and trafficking, providing a foundation for understanding and harnessing phage-host interfaces in human health and therapy.

## Materials and Methods

### Bacterial strains, bacteriophages, plasmids and media

For the cloning experiments, we used *E. cloni* 10G ELITE electrocompetent cells (Lucigen, 60080-2). *E. coli* EV36, an *E. coli* K-12-K1 hybrid derivative with the ability to express a Kl polysaccharide capsule morphologically similar to that of E. coli K1 clinical isolates^1^. The K1F phage, pSBC3 and pCas9K1FC2 plasmids were kind gifts of Tamás Fehér.

### Tissue cell culture

Three different epithelial cell lines were investigated: A549 (human tumorigenic lung epithelial cell line) (ATCC, CCL-185), Caco-2 (non- or low mucus-producing human colorectal adenocarcinoma cell line) (Sigma, 86010202) and HT29-MTX-E12 (mucus-producing human colon cancer cell line), which was derived from HT29 cells differentiated into mature goblet cells using methotrexate (Sigma, 12040401).

Unless otherwise indicated, all cell lines were cultured in Dulbecco′s Modified Eagle′s Medium (DMEM GlutaMAX) (Gibco, 21885-025) complemented with 10% fetal bovine serum (FBS) (Sigma, F4135), incubated at 37 °C in an incubator with an atmosphere containing 5% CO_2_, 95% relative humidity and grown to reach 80-90% confluency. To prevent bacterial contamination, the medium was supplemented with a 1x penicillin-streptomycin antibiotic mixture (Capricorn Scientific, PS-B). Before conducting any measurements, the medium was replaced with antibiotic-free medium. Time-lapse imaging of all cell lines was carried out in FluoroBrite DMEM Medium (Gibco, A1896701) supplemented with 10% FBS, while the microscopy measurements for detecting the colocalization pattern of phages with cell’s endomembranes were conducted in Ringer-Hepes solution (150 mM NaCl, 5.2 mM KCl, 2.2 mM CaCl2, 0.2 mM MgCl2, 1.2 mM MgSO4, 6 mM NaHCO3, 5 mM D-glucose, 10 mM Hepes, pH 7,4) supplemented with 1% FBS.

### Selection for epithelial adherent phages

To identify phages capable of adhering to the gut epithelial cells, first, we obtained a pooled fecal viral filtrate from fecal samples acquired from 10 healthy and 10 dysbiotic individuals ensuring high taxonomic phage diversity to increase the odds of recovering adherent phages. Fecal filtrates were prepared following a previously described protocol^2^. Briefly, 0.5 g of each fecal sample was placed in 10 mL of SM buffer (50 mM Tris-HCl, 100 mM NaCl, 8.5 mM MgSO4, pH 7.5), followed by vigorous vortexing for 5 minutes. The samples were then placed on ice for 5 minutes and centrifuged (5000 rpm, 10 minutes, 4°C). The supernatant was transferred to new Falcon tubes, and the centrifugation step was repeated. Finally, the supernatant containing the phage particles was filtered through a 0.45 μm pore size PES membrane. The filtrates were stored in a refrigerator at 4°C until use. As a positive control for adherence to the mucosal cell surface, we spiked the filtrate with T4 phage in 10^6^ PFU/mL final concentration per sample, similarly to how the spike-in control was used in a viral metagenomic study^3^.

Throughout the entire study, we complied with all relevant ethical regulations. The protocol related to human faecal sample collection was approved by the Ethical Review Board of the Albert Szent-Györgyi Health Centre, University of Szeged (approval ID: 42/2017-SZTE). Faeces were collected from 20 consenting Irish adult volunteers according to study protocol APC055, approved by the Cork Research Ethics Committee. Written informed consent from each participant was obtained before faecal sample collection.

To assess the adherence of the phages to the epithelial cell surface, the filtrate was applied on the confluent cell layer of two different gut epithelial cell types: Caco-2, a non- or low mucus-producing and HT29-MTX a mucus-producing cell line. Samples were incubated for 2-3 hrs at 37 °C in an incubator with an atmosphere of 5% CO_2_ and 95% relative humidity. Following incubation, the effluent fraction was collected by gentle removal of the supernatant. Cells were washed twice with 10 mL PBS and the liquid resulting from the first washing step was added to the supernatant containing the effluent phages. Following washing, adherent phages were eluted from the surface of the cell layer using an established methodology^3-5^. Briefly, cells were treated with 1,5 mL of Trypsin-EDTA to detach them from the culture dish, swayed gently for 1 min and incubated at 37 °C for 10 min (HT29-MTX cells) or 15 min (Caco-2 cells), respectively, accounting for the sensitivity of the cell lines against Trypsin-EDTA. As a next step, we added 10 mL of DMEM complemented with 10% of fetal bovine serum (FBS) to the samples. Cells were detached by a single shaking followed by their collection in Falcon tubes and their subsequent treatment with dithiothreitol (DTT) (20 mM final concentration). As DTT reduces disulfide bonds in mucins^6^, it was used to disassemble the adherent phages containing mucus layer from the cell surface. DTT-treated samples were incubated at 37 °C for 30 min, then kept on ice for 15 min and finally centrifuged at 4 °C for 20 min (4500-5000 rpm). The resulting supernatant was filtered by using a PES membrane with 0.45 μM pore diameter and samples were stored at 4 °C. The DNA extracted from these samples was sent for sequencing and the resulting data was subjected to in-depth bioinformatic analysis. To ensure reproducibility, the above-mentioned entire procedure was carried out in two biological replicates.

### Sequencing the genomic DNA from the viral filtrates

Phage genomic DNA was extracted from both the effluent and the residual fractions, as well as from the initial viral filtrate, and prepared for next-generation sequencing using the Accel-NGS 1S Plus DNA Library Kit (Swift Biosciences, 10024) following the manufacturer’s instructions. Samples were indexed by using the xGen™ CDI Primers primer package (Swift Biosciences; 10009794). The genomic DNA of the selected engineered phages were extracted using Norgen phage DNA extraction kit (Cat.n.: 46800, Norgen Biotek) and subsequently sequenced by Illumina shotgun NGS approach.

### Bioinformatic analysis of sequencing data

Assembly of reads was done using Metaphage 0.3.2^7^, a comprehensive viral assembly pipeline employing fastP and seqScreener for quality checking, Kraken2 and Krona for filtering non-viral reads, and Megahit for assembly, which is later evaluated by QUAST. Assembled contigs were passed to four viral screening tools, VIBRANT^8^, VirSorter^9^, Phigaro^10^ and VirFinder^11^ to distinguish viral elements (contigs of phage origin). The resulting 97460 contigs of phage origin were binned at 95% nucleotide sequence identity into vOTUs (Viral Operational Taxonomic Units), from which read count was calculated by BowTie2^12^ and consequently BamtoCov^13^. In all samples, more than 90% of reads passed quality filtering. Most viral elements were found by Vibrant or Virfinder, with Phigaro and Virsorter having a very low number (0-10) of hits exclusive to them. The largest contigs in each sample were always between 100 and 500000 base pairs, corresponding to the estimated scales of phage genome size (Table S1).

To remove redundant or duplicate sequences, vOTUs were dereplicated by MetaPhage^7^ using CD-HIT-EST^14^ with the default sequence identity threshold of 0.95 (95%), word size of 9, and alignment coverage of 0.85, according to the original paper. In some cases, when full or partial reverses of the same (or very similar) phage sequence occurred, two vOTUs were separated by the algorithm even though they shared a high (>97%) average nucleotide identity (ANI). In such instances, to demultiplex the data, we prioritized keeping the vOTU with the longer contigs. In cases where contig lengths were equal, preference was given to the one with a higher total read count.

To further ensure the high quality of the investigated data, the 3809 non-redundant vOTUs obtained after binning and dereplication were checked for genome completeness. vOTUs with less than 90% predicted genome completeness, calculated by CheckV v1.5^15^, were removed, resulting in 451 vOTUs. Finally, as a per-sample quality control, vOTUs were deemed present in a given sample if their corresponding coverage exceeded 1, otherwise they were considered absent (Figure 1B). In addition, vOTUs exclusively identified in the initial fecal filtrate samples were also excluded from the analysis.

### Classification of the vOTUs based on their adherence

After performing the above-elaborated quality filtering steps, 200 vOTUs that were retained (Table S2) were grouped into two categories: ‘adherent’ and ‘non-adherent’ vOTUs. For this categorization, as a first step, we determined the abundance of each vOTUs in different samples. To avoid overestimating the abundance of phages with larger genomes, we calculated their abundance by using the Lander-Waterman formula^16^, which takes into account the genome sizes of vOTUs relative to their read counts:

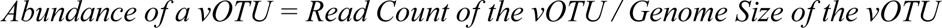

The resulting database was then used to define if a vOTU could be considered as adherent or non-adherent. For this, we employed two different statistical approaches, one that relied on the presence of vOTUs in the residual samples and one that measured vOTU depletion from the effluent samples.

We defined a vOTU as highly prevalent in residual samples if its abundance value, as calculated above, was higher than 1 in each of the residual replicates as well as was present in at least one of the original fecal filtrates to ensure it was not a contaminant. To estimate the level of depletion due to cellular uptake for each vOTU, we calculated the fold change between the ratio of a given vOTU in the initial filtrate and the ratio of a given vOTU in the effluent sample. For this, we applied the following formula:

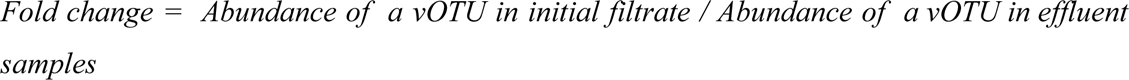

In cases when the resulting fold change value was higher than 1, meaning that the relative abundance in the initial filtrate was higher than in the effluent sample, we assumed that the phage was depleted from the effluent sample due to its adherence to the mucus and consequent internalization (Table S2).

In sum, a vOTU was considered adherent (‘adherent’ vOTU) if it met two criteria: i) it was clearly present (coverage > 1) in all four residual samples (two replicates with both HT29-MTX and Caco-2 cell lines), and ii) its abundance fold change between the initial filtrate and the effluent sample exceeded 1 across all four replicates, indicating consistent depletion from the effluent samples. Seventeen vOTUs met these criteria, while the remaining 200 were categorized as non-adherent (‘non-adherent’ vOTU) (Figure 1B and Table S2). As expected, the T4 phage, included as a positive control, was classified as adherent.

### Predicting the ORF- and domain content of vOTUs

To identify those genetic factors that could potentially confer phages with the ability to adhere to mucosal cell surfaces, the subset of 200 vOTUs was further investigated by predicting their ORF- and domain content. Ig-like-domain-containing proteins were our primary candidates for investigation in the context of adherence. Predictions were done by annotating all vOTUs with Pharokka^17^ and selecting those that had a domain of interest within 5 ORFs distance before or after the major or minor capsid proteins. Domain presence was determined by screening with Interpro’s Interproscan^18,19^ (release 5.65-97.0) standalone module for Immunoglobulin-like fold (IPR013783) and BACON (Bacteroidetes-Associated Carbohydrate-binding Often N-terminal) (IPR024361) domains.

### Estimating the abundance and prevalence of phages harbouring adherence-related domains in the human virome

To investigate whether mucosal surface binding is a common and broadly utilized feature of the human virome or whether phages carrying the experimentally characterized Ig-like domains are only exceptional cases, we analysed two datasets: (i) Cenote Human Virome Database^20^, encompassing seven human body sites from >6,000 metagenomes, comprising >45,000 vOTUs, designated here as the Broad Human Virome; and (ii) faecal viromes of 40 healthy individuals and 39 patients with Crohn’s disease or ulcerative colitis, consisted of >28,000 vOTUs^21^, designated here as 79-Individual Gut Virome (Tables S13 and S14, provided as separate Supplementary files).

By using CheckV we evaluated the completeness of these genomes and kept only the ones with higher than 90% completeness to avoid analysing genomes with a significant portion of their structural proteins likely missing. ORFs were retrieved by using a customary Python script, while domains were annotated with Interpro. We screened all phages for surface domains that were related to adherence, looking for matches with at least one of the following keywords: adherence, adhesin, adhesion, adsorption, bacon, big_, cadherin, carbohydrate-binding, cell-adhesion, cell-cell, cell-surface, concanavalin *-like, Fib_alpha, Fibronectin type III, fn2, fn3, glycan-binding, ig domain, ig like, Ig_J_chain, ig-fold, ig-like, immunoglobulin like, immunoglobulin-like, Importin_rep_3, intimin, invasin, invasion, I-set, lectin, leucine rich repeat, lrr, mucin, mucosal, mucus, peptidoglycan-binding, Phage_glycop_gL, pkd, protein-protein, sialidase, v-set. We used an E Value of 0.01 as a cutoff point.

Using diamond blastp^22^ the ORFs retrieved from both datasets were checked against the 45 ORFs detected in the 13 experimentally identified adherent vOTUs and those that shared ≥80% sequence identity and had a domains’ coverage ≥80% (475 in total) were clustered using Levenshtein distances^23^ with a similarity threshold of 80% (Tables S15 and S16).

To ensure unbiased comparisons, we restricted the analysis to high-quality phage genomes (>90% completeness and high or medium confidence), yielding 15556 and 2269 vOTUs from the Broad Human Virome and 79-Individual Gut Virome datasets, respectively. For both datasets, we performed enrichment analyses using Fisher’s exact test to assess whether Ig-like domain-encoding phages are enriched among highly abundant and highly prevalent phages compared with phages that do not encode these domains. For vOTUs belonging to the Broad Human Virome dataset, the enrichment analysis was performed across different body sites. For this, abundance and prevalence values were split into three categories (tertiles) and labeled as low, medium and high. For the Broad Human Virome dataset this was done for each body site separately. Enrichment analyses were performed only for the high abundance and high prevalence categories (Tables S17, S18 and S19).

To check whether vOTUs carrying at least one BACON, PKD, or unclassified Ig-like domain were associated with disease-associated microbiomes, for the Broad Human Virome dataset, we performed an enrichment analysis after grouping vOTUs into two categories based on whether they were significantly associated with disease status^20^ (Table S20). For the 79-Individual Gut Virome dataset we compared the abundance of Ig-like- domain-encoding phages between the healthy (n = 40) and the IBD (n = 39) cohorts by using Mann-Whitney U test (Table S14).

### Comparative genomics map

To visualize the taxonomic relationships of the identified vOTUs, we used the classical approach of constructing an orthogroup-based network of viruses, where proteins of all participant phages are clustered, and the phages’ level of connection is determined by the number of proteins they have in common viral clusters (VCs).

The network was calculated by Metaphage, applying vConTACT2’s^24^ gene2genome module, an orthogroup-based classifier, that clusters proteins into homologous orthogroups, so-called viral clusters. The reference database used in vConTACT2 was provided by Metaphage. The connectivity of nodes representing our phages is determined by the percentage of orthogroups they share. The visualization of the vConTACT2 network was performed by using Cytoscape^25^.

### Engineering of K1F phages displaying Ig-like-domain-containing proteins

K1F phages displaying the identified Ig-like-domain-containing proteins were constructed by using homology-based recombination coupled with CRISPR/Cas9 counterselection method described in a previous study^26^, with minor modifications. We used the non-adherent phage K1F as a scaffold to display a selected set of Ig-like domain-containing proteins identified as a result of our screen for adherent phages. For candidate domains, we selected eight Ig-like domain-containing proteins from different clusters, which were carried by phages defined as adherent. The corresponding genes were acquired by PCR amplification from the initial fecal viral filtrate and 3 Ig-like domain-containing protein-coding genes were successfully integrated into the 3’ end of the minor capsid gene of the K1F phage just before the stop codon, alongside a linker region (Figure S2). In addition to the 3 newly identified Ig-like domain-containing proteins, we engineered a K1F variant that displays the Hoc protein of the T4 phage. We used this construct as a positive control for adherence, as this protein was identified as the key determinant by which T4 adheres to mucus residues and also carries three Ig-like domains^27^.

In brief, wild-type K1F phages were propagated on *E. coli* EV36 host strain carrying the pSBC3 donor plasmid featuring the gene of interest flanked by sequences homologous with the targeted region. The resulting lysates were mixtures of wild-type and recombinant phages, predominated by the former. To increase the ratio of recombinants, the resulting mixture of phages was next propagated on the EV36 host harboring an CRISPR/Cas9 and guide RNA containing plasmid. We used this plasmid to target the 3’ end of the unmodified minor capsid gene sequence of the K1F, thus selecting against the wild-type phages (for details, see Figure S2). After performing a plaque assay with the acquired phage lysates, the recombinants were identified from the resulting unique plaques with PCR using the appropriate primers (Table S13). As a final confirmation of the integration, the genomic DNA of each engineered phage were extracted using a Norgen phage DNA extraction kit (Norgen Biotek) and sequenced by Illumina shotgun NGS approach.

### Phage propagation and titering

For phage propagation we used the *E. coli* EV36 strain. Overnight bacterial culture was prepared from glycerol stock by inoculation into 3 mL LB medium in a 13 mL tube and incubated at 37°C with shaking at 220 rpm. The following day, the overnight culture was diluted in a 1:100 ratio in fresh LB medium and incubated at 37°C with shaking at 220 rpm until the optical density (OD_600_) reached 0.6, after which it was inoculated with the phage stock at a multiplicity of infection (MOI) of 0.01. The phage-infected culture was then incubated at 37°C with shaking until complete bacterial lysis was observed, indicated by the clearance of the culture (typically after 2-3 hours of incubation). After observing complete lysis, the culture was transferred to sterile centrifuge tubes, chilled on ice for 15 minutes and centrifuged at 4500 rpm for 10 minutes at 5°C to pellet any remaining bacterial debris. To obtain a phage lysate devoid of bacterial cells, the supernatant was filtered twice: first through a 0.45 µm PES syringe filter, followed by filtration through a 0.25 µm PES syringe filter (VWR, 76479-026). Finally, the lysate was stored at 4°C.

The titer of the phage lysates was determined by using the classic plaque assay protocol. 400 µl of mid-log-phase (OD_600_ ≈ 0.6) *E. coli* EV36 culture was added to 4 mL of molten top agar (LB supplemented with 0.7% agar, kept at approximately 45°C), mixed gently, poured evenly on the surface of a pre-warmed LB agar plate and allow the top agar to solidify at room temperature. Serial 10-fold dilutions (ranging from 10^-1^ to 10^-9^) of the phage lysate were prepared in LB medium and from each dilution 10 µL was pipetted in duplicate onto the surface of the plates prepared as described above. After the droplets dried, the plates were incubated inverted overnight at 37 °C and the following day the number of plaques was counted to determine the phage titer expressed in PFU/mL.

### Live cell imaging

For the live cell imaging, cells were seeded in Greiner µClear black 96-well plates (Greiner, 655090). Before microscopy, the antibiotic-containing culture medium was replaced with antibiotic-free medium supplemented with Hoechst dye (1 µg/mL final concentration)(Sigma, B2261) and cells were incubated for 20 minutes (37 °C, 5% CO_2_). After the incubation, cells were washed twice with PBS solution and then kept in antibiotic-free medium until further use.

Phages used in microscopy experiments were labeled with 1% SYBR™ Gold nucleic acid dye (Thermo Fisher Scientific, S11494), following the protocol described by Bichet and colleagues^28^. Time-lapse image sequences of A549 and HT29MTX cell lines were obtained using the Operetta CLS high content analysis system (PerkinElmer) with a 40x water objective (NA 1.1) in confocal mode and time-lapse image sequences of Caco2 cell line were obtained using the Operetta high-content screening system (PerkinElmer) with a 60x air objective (NA 0.7) in widefield mode. Scanning was performed using a brightfield channel to identify cells, while fluorescent channels; DAPI (355/385, 430/500), EGFP (430/475, 525/550) were used to detect intracellular signals of the fluorescent dyes. For each field of view, a 6-frame image sequence was acquired with a 4-hour gap between subsequent frames. 140 fields of view were acquired for each slide using six (A549, HT29MTX) and one (Caco2) Z focus plane determined by the laser-based autofocus system.

The phage-endomembrane colocalization experiments were implemented using Operetta CLS high content analysis system with a 40x water objective (NA 1.1) Pictures were taken in confocal mode with 12 stacks using brightfield channel to identify cells, while fluorescent channels DAPI (355/385, 430/500), EGFP (460/490, 500/550), mCherry (530/560, 570/650) and Deep Red (615/645, 655/760) were used to detect intracellular signals of the fluorescent dyes. All of Z-stacks were taken across the entire nuclear depth, from the apical to the basal height.

### Image analysis pipeline of phage uptake

Screens performed by Operetta were processed using BIAS (BioImage Analysis Software)^29^. During the pre-processing step, non-uniform illumination was corrected separately for each channel by using the CIDRE method^30^. The nucleAIzer basic nucleus segmentation model^31^ integrated into BIAS was applied to detect individual nuclei in the images. In segmentation post-processing, two additional regions were defined for each nucleus: a) a region representing the entire cell was defined by extending nuclei regions up to a maximum radius of 10 μm so that adjacent cells did not overlap, b) cytoplasmic regions were defined by subtracting nuclei segmentation masks from extended cell masks. Finally, morphological properties of these three different regions as well as intensity and texture features from all channels were extracted (in total 228 features) for cell classification. We employed supervised machine learning to predict three different cell types: GFP-positive cells, GFP-negative cells and other cells or segmentation artefacts that can be considered trash. These classes were manually selected based on their morphological characteristics.

The data obtained from the microscopy experiments were processed as follows: For each sample, at each timepoint 27 microscope field-of-views (FOVs) were analysed. For each FOV, the percentage of cells with internalized phages, representing the phage uptake by the epithelial cells, was calculated from the ratio of cells with internalized phages observed in a microscope field of view. This process was also applied to the medium control sample, where no phages were added to the cells. For this control sample, the median percentage value of cells with internalized phages was calculated from 27 FOVs at each timepoint. Following this, for each sample, for each FOV per timepoint, the percentage of cells with internalized phages was normalized by subtracting the median percentage value of the control sample. If the resulting value was negative, it was set to zero (see Table S9). Area under curve (AUC) representation of the phage uptake was calculated from the normalized percentages of cells with intracellular phages and samples were compared by using Kruskal-Wallis test.

To compare the fold change decrease in uptake of the wild-type and engineered K1F phages, and the T4 phage by HT29-MTX epithelial cells in response to different treatments of the epithelial cell surface, a fold change value was calculated based on the following formula: fold change = median AUC in the case of untreated cells / AUC for each FOV in the case of treated cells. This was performed separately for each of the three treatments: N-acetylcysteine (NAC), cyclosporin A (CSA) and heparinase I (HEP)(Table S10).

### Identification of interaction modes of Ig-like domain-carrying engineered phages

To determine the possible binding sites of our engineered K1F phages, HT29-MTX cells were subjected to the following different treatments: i) N-acetyl-L-cysteine treatment, to degrade the mucin crosslinking disulfide bonds; ii) Cyclosporine A treatment, for mucin aggregation and heparan sulfate shedding, and iii) heparinase I treatment to cleave highly sulfated heparin/HS chains.

A monolayer of HT29-MTX cells grown in Greiner black 96-well plates (Greiner, 655090) was treated for 1 hour at 37°C either with 15 mM N-acetyl-L-cysteine or 1 mM Cyclosporine A. In the case of heparinase I, cells were treated for 2 hrs at 37°C with 3 U/mL enzymes. After the treatment, cells were washed twice with PBS buffer, the labeled phages were added in 10^9^ PFU/mL concentration and the phage uptake was investigated with the Operetta high content analysis system after 18 hours. Control cells did not receive any treatment.

### Determining the intracellular fates of internalized phages

To track phages after their internalization by the human epithelial cells and determine their intracellular fates, we used live-cell markers to check the co-localization of the stained phages with different labeled cell organelles. To specifically detect the membranes of mature lysosomes, endoplasmatic reticulum and Golgi apparatus, we performed live-cell staining with cell-permeant Lysotracker^TM^ Deep Red (Invitrogen, L12492), ER-Tracker^TM^ Red (Invitrogen, E34250) and BODIPY^TM^ TR Ceramide (Invitrogen, D7540) dyes, respectively. We followed the protocol described by the manufacturer with slight modifications. In brief, SYBR-Gold-stained phages were applied in 10^8^ PFU concentration onto A549 epithelial cells and incubated for 6 and 18 hours. After incubation, phages were removed, and the samples were washed three times with PBS.

To visualize the lysosomes and ER, the corresponding dye was added to the cells in a fresh antibiotic-free medium at a final concentration of 1 μM and incubated for 30 minutes. After washing the cells three times with PBS, Ringer-Hepes solution supplemented with 1% FBS was added. To visualize the Golgi apparatus, the corresponding dye was added to the cells in 3% BSA containing PBS solution at a final concentration of 5μM, followed by incubation at 4 °C for 30 min. Cells were washed three times with ice-cold antibiotic-free medium, fresh medium added and and incubated at 37 °C for 30min. After washing three times with PBS, Ringer-Hepes solution supplemented with 1% FBS was added. Colocalization analysis was performed on paired fluorescence images by extracting per-pixel red (Lysotracker^TM^ Deep Red, ER-Tracker^TM^ Red or BODIPY^TM^ TR Ceramide) and SYBR Gold (green) channel intensities. After background thresholding, Manders’ overlap coefficients (M1 and M2) and conditional overlap fractions were calculated based on intensity positive pixels (Table S22).

### Live cell confocal microscopy

The images of the live cells were taken by Leica Stellaris 5 confocal laser scanning microscope (CLSM) with culture chamber (to maintain 37 °C and 5% CO_2_) and a HC PL APO CS2 63x/1.40 oil immersion objective. We used hybrid detectors (HyD) to visualise nucleus (Hoechst 33342, Sigma) (HyD X 2, excitation/emission 361/497 nm), phage DNA (SYBR Gold, Thermo Scientific) (HyD X 2 excitation/emission 495/537 nm), endoplasmatic reticulum (ER-Tracker Red) (Invitrogen) (HyD R 5, excitation/emission 587/615), golgi apparatus BODIPY TR Ceramide (Invitrogen) (HyD R 5, excitation/emission 589/617)

### Monitoring phage shedding in mouse feces

To compare the fecal shedding kinetics of the engineered K1F phages with that of the wild-type K1F phage and T4 phage, 8-10 week-old (18-20 g) female BALB/c OlaHsd mice were used. Animals were cared for in accordance with the guidelines of the European Federation for Laboratory Animal Science Associations (FELASA) and all procedures, care and handling of the animals were approved by the Animal Welfare Committee of the Enviroinvest Co., Pécs (Permit Number: BA02/2000-12/2022). For the experiment, animals were divided into six groups, but each animal was housed in a separate cage. Bacteriophages were administered without prior treatment or food deprivation. Each animal received 200 µl of synchronised phage suspension (10^8^ PFU/mL) through a special gastric tube. Stool samples (3-4 pieces, equalling 50-100 mg) were collected from the animals 2h, 4h, 6h, 8h, 10h, 12h, 16h, 20h and 24h after phage administration and were thoroughly suspended in PBS by pipetting and vortexing. The so-gained suspensions were centrifuged (10.000xg, 4min) and 200 µl supernatants were transferred to new reagent tubes. Supernatants were treated with chloroform in a 1:50 ratio (4 µl) and incubated in the fridge for a couple of hours. After incubation, PFU was determined by diluting the supernatants 10-fold (-1, -2, -3) in PBS and pipetting 10-10 µl both from the diluted and undiluted samples onto the surface of the plates containing freshly spread bacterial lawn. Samples collected from animals treated with K1F phages (K1F, K1F::S, K1F::L, K1F::7649 and K1F::hoc) were tested on the E. coli strain EV36, while those treated with T4 phage on *E. coli* strain BW25113.

## Data availability

Sequencing reads generated in this study have been deposited in the European Nucleotide Archive (ENA) under the study accession PRJEB107177. All other data supporting the findings of this study are available within the paper and its Supplementary Information or from the corresponding authors upon request.

## Acknowledgement

This work was supported by the National Research, Development and Innovation Fund ADVANCED 149516 grant, the 2024-1.2.2-ERA_NET-2024-00004 grant, the National Laboratory of Biotechnology grant 2022-2.1.1-NL-2022-00008, and the HUN-REN TKCS-2024/66 grant; the European Union’s Horizon 2020 Research and Innovation Programme no. 739593. Andrey Shkoporov was supported by the Wellcome Trust Research Career Development Fellowship [220646/Z/20/Z]. This research was funded in whole, or in part, by the Wellcome Trust [220646/Z/20/Z]. Gábor Apjok was supported by an EMBO Scientific Exchange Grant (Grant No. 9262). For the purpose of open access, the authors have applied a CC BY public copyright licence to any Author Accepted Manuscript version arising from this submission. We thank Tamás Fehér for providing the K1F bacteriophage display system, and Ferenc Kovács, András Kriston and Single-Cell Technologies Ltd. for providing access to the BIAS image analysis software.

## Author contributions

G.A. conceived the study and designed and performed experiments and bioinformatic analyses. T.S. designed and performed experiments and conducted image analysis. B.V. and A.A. contributed to bioinformatic analyses. D.S., L.B. and I.G. contributed to experimental work. P.H., S.J. and E.M. performed microscopy and image analysis. I.G. and M.D. contributed to human cell line cultivation. G.S. carried out mouse experiments. A.S. and C.H. contributed to virome analysis and interpretation. B.K., C.P. and A.S. provided resources and supervision. G.A., B.K. and O.M. wrote the manuscript with input from all authors.

## Correspondence

Correspondence and requests for materials should be addressed to G.A. or B.K.

## Supplementary Information

Supplementary Information is available for this paper.

## Reference

1. Mirzaei, M. K. & Maurice, C. F. Ménage à trois in the human gut: interactions between host, bacteria and phages. Nat. Rev. Microbiol. 15, 397–408 (2017).

2. Ma, Z., Zuo, T., Frey, N. & Rangrez, A. Y. A systematic framework for understanding the microbiome in human health and disease: from basic principles to clinical translation. Signal Transduct. Target. Ther. 9, 237 (2024).

3. Zhang, Y. & Wang, R. The human gut phageome: composition, development, and alterations in disease. Front. Microbiol. 14, (2023).

4. Bodner, K., Melkonian, A. L. & Covert, M. W. The Enemy of My Enemy: New Insights Regarding Bacteriophage-Mammalian Cell Interactions. Trends Microbiol. 29, 528–541 (2021).

5. Barr, J. J. A bacteriophages journey through the human body. Immunol. Rev. 279, 106–122 (2017).

6. Haddock, N. L. et al. Phage diversity in cell-free DNA identifies bacterial pathogens in human sepsis cases. Nat. Microbiol. 8, 1495–1507 (2023).

7. Shkoporov, A. N. et al. Viral biogeography of the mammalian gut and parenchymal organs. Nat. Microbiol. 7, 1301–1311 (2022).

8. Maľarik, T., Bhide, K., Talpašová, L. & Bhide, M. Mphages and the Blood-Brain Barrier: A Review. Folia Vet. 68, 15–21 (2024).

9. Van Belleghem, J. D., Dąbrowska, K., Vaneechoutte, M., Barr, J. J. & Bollyky, P. L. Interactions between Bacteriophage, Bacteria, and the Mammalian Immune System. Viruses 11, 10 (2018).

10. Hibstu, Z., Belew, H., Akelew, Y. & Mengist, H. M. Phage Therapy: A Different Approach to Fight Bacterial Infections. Biol. Targets Ther. 16, 173–186 (2022).

11. Roach, D. R. et al. Synergy between the Host Immune System and Bacteriophage Is Essential for Successful Phage Therapy against an Acute Respiratory Pathogen. Cell Host Microbe 22, 38–47.e4 (2017).

12. Bichet, M. C. et al. Mammalian cells internalize bacteriophages and use them as a resource to enhance cellular growth and survival. PLOS Biol. 21, e3002341 (2023).

13. Manrique, P. et al. Healthy human gut phageome. Proc. Natl. Acad. Sci. U. S. A. 113, 10400–10405 (2016).

14. Adiliaghdam, F., et al. Human enteric viruses autonomously shape inflammatory bowel disease phenotype through divergent innate immunomodulation. Sci. Immunol. 7, eabn6660 (2022).

15. Mayneris-Perxachs, J. et al. Caudovirales bacteriophages are associated with improved executive function and memory in flies, mice, and humans. Cell Host Microbe 30, 340–356.e8 (2022).

16. Darfeuille-Michaud, A. et al. High prevalence of adherent-invasive Escherichia coli associated with ileal mucosa in Crohn’s disease. Gastroenterology 127, 412–421 (2004).

17. Moussata, D. et al. Confocal laser endomicroscopy is a new imaging modality for recognition of intramucosal bacteria in inflammatory bowel disease in vivo. Gut 60, 26–33 (2011).

18. Ogawa, M. & Sasakawa, C. Intracellular survival of Shigella. Cell. Microbiol. 8, 177–184 (2006).

19. Palmela, C. et al. Adherent-invasive Escherichia coli in inflammatory bowel disease. Gut 67, 574–587 (2018).

20. Pickard, J. M., Zeng, M. Y., Caruso, R. & Núñez, G. Gut microbiota: Role in pathogen colonization, immune responses, and inflammatory disease. Immunol. Rev. 279, 70–89 (2017).

21. Josenhans, C., Müthing, J., Elling, L., Bartfeld, S. & Schmidt, H. How bacterial pathogens of the gastrointestinal tract use the mucosal glyco-code to harness mucus and microbiota: New ways to study an ancient bag of tricks. Int. J. Med. Microbiol. 310, 151392 (2020).

22. Nguyen, S. et al. Bacteriophage Transcytosis Provides a Mechanism To Cross Epithelial Cell Layers. mBio 8, e01874–17 (2017).

23. Rothschild-Rodriguez, D., Hedges, M., Kaplan, M., Karav, S. & Nobrega, F. L. Phage-encoded carbohydrate-interacting proteins in the human gut. Front. Microbiol. 13, 1083208 (2023).

24. Almeida, G. M. F., Laanto, E., Ashrafi, R. & Sundberg, L.-R. Bacteriophage Adherence to Mucus Mediates Preventive Protection against Pathogenic Bacteria. mBio 10, e01984–19 (2019).

25. Fu, K. et al. Escherichia coli phage ΦPNJ-9 adheres to mucus via a variant Hoc protein. J. Virol. 99, e0178924 (2025).

26. Fraser, J. S., Yu, Z., Maxwell, K. L. & Davidson, A. R. Ig-Like Domains on Bacteriophages: A Tale of Promiscuity and Deceit. J. Mol. Biol. 359, 496–507 (2006).

27. Bayfield, O. W. et al. Structural atlas of a human gut crassvirus. Nature 617, 409–416 (2023).

28. Barr, J. J. et al. Bacteriophage adhering to mucus provide a non-host-derived immunity. Proc. Natl. Acad. Sci. 110, 10771–10776 (2013).

29. Bichet, M. C. et al. Bacteriophage uptake by mammalian cell layers represents a potential sink that may impact phage therapy. iScience 24, 102287 (2021).

30. Rao, V. B. & Zhu, J. Bacteriophage T4 as a nanovehicle for delivery of genes and therapeutics into human cells. Curr. Opin. Virol. 55, 101255 (2022).

31. Kan, L. & Barr, J. J. A Mammalian Cell’s Guide on How to Process a Bacteriophage. Annu. Rev. Virol. 10, 183–198 (2023).

32. Tobin, C. A., Hill, C. & Shkoporov, A. N. Factors Affecting Variation of the Human Gut Phageome. Annu. Rev. Microbiol. 77, 363–379 (2023).

33. Majzoub, M. E. et al. The phageome of patients with ulcerative colitis treated with donor fecal microbiota reveals markers associated with disease remission. Nat. Commun. 15, 8979 (2024).

34. Yu, X. et al. Gut phageome: challenges in research and impact on human microbiota. Front. Microbiol. 15, (2024).

35. Gagnon, M., Zihler Berner, A., Chervet, N., Chassard, C. & Lacroix, C. Comparison of the Caco-2, HT-29 and the mucus-secreting HT29-MTX intestinal cell models to investigate Salmonella adhesion and invasion. J. Microbiol. Methods 94, 274-279 (2013).

36. Chin, W. H. et al. Bacteriophages evolve enhanced persistence to a mucosal surface. Proc. Natl. Acad. Sci. U. S. A. 119, e2116197119 (2022).

37. Hughes, J. et al. The polycystic kidney disease 1 (PKD1) gene encodes a novel protein with multiple cell recognition domains. Nat. Genet. 10, 151–160 (1995).

38. Møller-Olsen, C., Ho, S. F. S., Shukla, R. D., Feher, T. & Sagona, A. P. Engineered K1F bacteriophages kill intracellular Escherichia coli K1 in human epithelial cells. Sci. Rep. 8, 17559 (2018).

39. Martínez-Maqueda, D., Miralles, B. & Recio, I. HT29 Cell Line. in The Impact of Food Bioactives on Health: in vitro and ex vivo models (eds Verhoeckx, K., et al.) (Springer, Cham (CH), 2015).

40. Horibe, S., Tanahashi, T., Kawauchi, S., Murakami, Y. & Rikitake, Y. Mechanism of recipient cell-dependent differences in exosome uptake. BMC Cancer 18, 47 (2018).

41. Fokine, A. et al. Structure of the three N-terminal immunoglobulin domains of the highly immunogenic outer capsid protein from a T4-like bacteriophage. J. Virol. 85, 8141–8148 (2011).

42. Scholl, D. & Merril, C. The genome of bacteriophage K1F, a T7-like phage that has acquired the ability to replicate on K1 strains of Escherichia coli. J. Bacteriol. 187, 8499–8503 (2005).

43. Kemp, P., Garcia, L. R. & Molineux, I. J. Changes in bacteriophage T7 virion structure at the initiation of infection. Virology 340, 307–317 (2005).

44. Sathaliyawala, T. et al. Functional analysis of the highly antigenic outer capsid protein, Hoc, a virus decoration protein from T4-like bacteriophages. Mol. Microbiol. 77, 444-455 (2010).

45. Weiss, M. et al. In vivo replication of T4 and T7 bacteriophages in germ-free mice colonized with Escherichia coli. Virology 393, 16–23 (2009).

46. Tisza, M. J. & Buck, C. B. A catalog of tens of thousands of viruses from human metagenomes reveals hidden associations with chronic diseases. Proc. Natl. Acad. Sci. 118, e2023202118 (2021).

47. Stockdale, S. R. et al. Interpersonal variability of the human gut virome confounds disease signal detection in IBD. Commun. Biol. 6, 221 (2023).

48. Tailford, L. E., Crost, E. H., Kavanaugh, D. & Juge, N. Mucin glycan foraging in the human gut microbiome. Front. Genet. 6, (2015).

49. Pedre, B., Barayeu, U., Ezeriņa, D. & Dick, T. P. The mechanism of action of N-acetylcysteine (NAC): The emerging role of H2S and sulfane sulfur species. Pharmacol. Ther. 228, 107916 (2021).

50. Kishimoto, H., Ridley, C., Inoue, K. & Thornton, D. J. Assessment of polymeric mucin-drug interactions. PLOS ONE 19, e0306058 (2024).

51. Kishimoto, H., Ridley, C. & Thornton, D. J. The lipophilic cyclic peptide cyclosporin A induces aggregation of gel-forming mucins. Sci. Rep. 12, 6153 (2022).

52. Kesimer, M. et al. Molecular organization of the mucins and glycocalyx underlying mucus transport over mucosal surfaces of the airways. Mucosal Immunol. 6, 379–392 (2013).

53. Florian, J. A. et al. Heparan sulfate proteoglycan is a mechanosensor on endothelial cells. Circ. Res. 93, e136–142 (2003).

54. Green, S. I. et al. Targeting of Mammalian Glycans Enhances Phage Predation in the Gastrointestinal Tract. mBio 12, e03474–20 (2021).

55. Kaksonen, M. & Roux, A. Mechanisms of clathrin-mediated endocytosis. Nat. Rev. Mol. Cell Biol. 19, 313–326 (2018).

56. Kiss, A. L. & Botos, E. Endocytosis via caveolae: alternative pathway with distinct cellular compartments to avoid lysosomal degradation? J. Cell. Mol. Med. 13, 1228–1237 (2009).

57. Mello, L. V., Chen, X. & Rigden, D. J. Mining metagenomic data for novel domains: BACON, a new carbohydrate-binding module. FEBS Lett. 584, 2421–2426 (2010).

58. Bycroft, M. et al. The structure of a PKD domain from polycystin-1: implications for polycystic kidney disease. EMBO J. 18, 297–305 (1999).

59. Mallard, F. et al. Direct pathway from early/recycling endosomes to the Golgi apparatus revealed through the study of shiga toxin B-fragment transport. J. Cell Biol. 143, 973–990 (1998).

60. Ribeiro, C. M. P. & Boucher, R. C. Role of endoplasmic reticulum stress in cystic fibrosis-related airway inflammatory responses. Proc. Am. Thorac. Soc. 7, 387–394 (2010).

61. M, L., E, R. & E, B. Impact of ER stress and the unfolded protein response on Fabry disease. EBioMedicine 115, (2025).

62. Yl, T. et al. ERdj3 is an endoplasmic reticulum degradation factor for mutant glucocerebrosidase variants linked to Gaucher’s disease. Chem. Biol. 21, (2014).

63. Vimr, E. R. & Troy, F. A. Regulation of sialic acid metabolism in Escherichia coli: role of N-acylneuraminate pyruvate-lyase. J. Bacteriol. 164, 854–860 (1985).

64. Shkoporov, A. N. et al. ΦCrAss001 represents the most abundant bacteriophage family in the human gut and infects Bacteroides intestinalis. Nat. Commun. 9, 4781 (2018).

65. Meldrum, O. W. et al. Mucin gel assembly is controlled by a collective action of non-mucin proteins, disulfide bridges, Ca2+-mediated links, and hydrogen bonding. Sci. Rep. 8, 5802 (2018).

66. Pandolfo, M., Telatin, A., Lazzari, G., Adriaenssens, E. M. & Vitulo, N. MetaPhage: an Automated Pipeline for Analyzing, Annotating, and Classifying Bacteriophages in Metagenomics Sequencing Data. mSystems 7, e0074122 (2022).

67. Kieft, K., Zhou, Z. & Anantharaman, K. VIBRANT: automated recovery, annotation and curation of microbial viruses, and evaluation of viral community function from genomic sequences. Microbiome 8, 90 (2020).

68. Guo, J. et al. VirSorter2: a multi-classifier, expert-guided approach to detect diverse DNA and RNA viruses. Microbiome 9, 37 (2021).

69. Phigaro: high-throughput prophage sequence annotation | Bioinformatics | Oxford Academic. https://academic.oup.com/bioinformatics/article/36/12/3882/5822875.

70. Ren, J., Ahlgren, N. A., Lu, Y. Y., Fuhrman, J. A. & Sun, F. VirFinder: a novel k-mer based tool for identifying viral sequences from assembled metagenomic data. Microbiome 5, 69 (2017).

71. Langmead, B. & Salzberg, S. L. Fast gapped-read alignment with Bowtie 2. Nat. Methods 9, 357–359 (2012).

72. Birolo, G. & Telatin, A. BamToCov: an efficient toolkit for sequence coverage calculations. Bioinformatics 38, 2617–2618 (2022).

73. Li, W. & Godzik, A. Cd-hit: a fast program for clustering and comparing large sets of protein or nucleotide sequences. Bioinformatics 22, 1658–1659 (2006).

74. Nayfach, S. et al. CheckV assesses the quality and completeness of metagenome-assembled viral genomes. Nat. Biotechnol. 39, 578–585 (2021).

75. Lander, E. S. & Waterman, M. S. Genomic mapping by fingerprinting random clones: a mathematical analysis. Genomics 2, 231–239 (1988).

76. Pharokka: a fast scalable bacteriophage annotation tool | Bioinformatics | Oxford Academic. https://academic.oup.com/bioinformatics/article/39/1/btac776/6858464.

77. Jones, P. et al. InterProScan 5: genome-scale protein function classification. Bioinformatics 30, 1236–1240 (2014).

78. Blum, M. et al. InterPro: the protein sequence classification resource in 2025. Nucleic Acids Res. 53, D444–D456 (2025).

79. Buchfink, B., Reuter, K. & Drost, H.-G. Sensitive protein alignments at tree-of-life scale using DIAMOND. Nat. Methods 18, 366–368 (2021).

80. Young, B., Faris, T. & Armogida, L. Levenshtein distance as a measure of accuracy and precision in forensic PCR-MPS methods. Forensic Sci. Int. Genet. 55, 102594 (2021).

81. Bin Jang, H., et al. Taxonomic assignment of uncultivated prokaryotic virus genomes is enabled by gene-sharing networks. Nat. Biotechnol. 37, 632–639 (2019).

82. Shannon, P. et al. Cytoscape: a software environment for integrated models of biomolecular interaction networks. Genome Res. 13, 2498–2504 (2003).

83. Wu, J. et al. Bacteriophage defends murine gut from Escherichia coli invasion via mucosal adherence. Nat. Commun. 15, 4764 (2024).

84. Mund, A. et al. Deep Visual Proteomics defines single-cell identity and heterogeneity. Nat. Biotechnol. 40, 1231–1240 (2022).

85. Smith, K. et al. CIDRE: an illumination-correction method for optical microscopy. Nat. Methods 12, 404–406 (2015).

86. Hollandi, R. et al. nucleAIzer: A Parameter-free Deep Learning Framework for Nucleus Segmentation Using Image Style Transfer. Cell Syst. 10, 453–458.e6 (2020).

## References

1. Vimr, E. R. & Troy, F. A. Regulation of sialic acid metabolism in Escherichia coli: role of N-acylneuraminate pyruvate-lyase. J. Bacteriol. 164, 854–860 (1985).

2. Shkoporov, A. N. et al. ΦCrAss001 represents the most abundant bacteriophage family in the human gut and infects Bacteroides intestinalis. Nat. Commun. 9, 4781 (2018).

3. Shkoporov, A. N. et al. Viral biogeography of the mammalian gut and parenchymal organs. Nat. Microbiol. 7, 1301–1311 (2022).

4. Tobin, C. A., Hill, C. & Shkoporov, A. N. Factors Affecting Variation of the Human Gut Phageome. Annu. Rev. Microbiol. 77, 363–379 (2023).

5. Majzoub, M. E. et al. The phageome of patients with ulcerative colitis treated with donor fecal microbiota reveals markers associated with disease remission. Nat. Commun. 15, 8979 (2024).

6. Meldrum, O. W. et al. Mucin gel assembly is controlled by a collective action of non-mucin proteins, disulfide bridges, Ca2+-mediated links, and hydrogen bonding. Sci. Rep. 8, 5802 (2018).

7. Pandolfo, M., Telatin, A., Lazzari, G., Adriaenssens, E. M. & Vitulo, N. MetaPhage: an Automated Pipeline for Analyzing, Annotating, and Classifying Bacteriophages in Metagenomics Sequencing Data. mSystems 7, e0074122 (2022).

8. Kieft, K., Zhou, Z. & Anantharaman, K. VIBRANT: automated recovery, annotation and curation of microbial viruses, and evaluation of viral community function from genomic sequences. Microbiome 8, 90 (2020).

9. Guo, J. et al. VirSorter2: a multi-classifier, expert-guided approach to detect diverse DNA and RNA viruses. Microbiome 9, 37 (2021).

10. Phigaro: high-throughput prophage sequence annotation | Bioinformatics | Oxford Academic. https://academic.oup.com/bioinformatics/article/36/12/3882/5822875.

11. Ren, J., Ahlgren, N. A., Lu, Y. Y., Fuhrman, J. A. & Sun, F. VirFinder: a novel k-mer based tool for identifying viral sequences from assembled metagenomic data. Microbiome 5, 69 (2017).

12. Langmead, B. & Salzberg, S. L. Fast gapped-read alignment with Bowtie 2. Nat. Methods 9, 357–359 (2012).

13. Birolo, G. & Telatin, A. BamToCov: an efficient toolkit for sequence coverage calculations. Bioinformatics 38, 2617–2618 (2022).

14. Li, W. & Godzik, A. Cd-hit: a fast program for clustering and comparing large sets of protein or nucleotide sequences. Bioinformatics 22, 1658–1659 (2006).

15. Nayfach, S. et al. CheckV assesses the quality and completeness of metagenome-assembled viral genomes. Nat. Biotechnol. 39, 578–585 (2021).

16. Lander, E. S. & Waterman, M. S. Genomic mapping by fingerprinting random clones: a mathematical analysis. Genomics 2, 231–239 (1988).

17. Pharokka: a fast scalable bacteriophage annotation tool | Bioinformatics | Oxford Academic. https://academic.oup.com/bioinformatics/article/39/1/btac776/6858464.

18. Jones, P. et al. InterProScan 5: genome-scale protein function classification. Bioinformatics 30, 1236–1240 (2014).

19. Blum, M. et al. InterPro: the protein sequence classification resource in 2025. Nucleic Acids Res. 53, D444–D456 (2025).

20. Tisza, M. J. & Buck, C. B. A catalog of tens of thousands of viruses from human metagenomes reveals hidden associations with chronic diseases. Proc. Natl. Acad. Sci. 118, e2023202118 (2021).

21. Stockdale, S. R. et al. Interpersonal variability of the human gut virome confounds disease signal detection in IBD. Commun. Biol. 6, 221 (2023).

22. Buchfink, B., Reuter, K. & Drost, H.-G. Sensitive protein alignments at tree-of-life scale using DIAMOND. Nat. Methods 18, 366–368 (2021).

23. Young, B., Faris, T. & Armogida, L. Levenshtein distance as a measure of accuracy and precision in forensic PCR-MPS methods. Forensic Sci. Int. Genet. 55, 102594 (2021).

24. Bin Jang, H., et al. Taxonomic assignment of uncultivated prokaryotic virus genomes is enabled by gene-sharing networks. Nat. Biotechnol. 37, 632–639 (2019).

25. Shannon, P. et al. Cytoscape: a software environment for integrated models of biomolecular interaction networks. Genome Res. 13, 2498–2504 (2003).

26. Møller-Olsen, C., Ho, S. F. S., Shukla, R. D., Feher, T. & Sagona, A. P. Engineered K1F bacteriophages kill intracellular Escherichia coli K1 in human epithelial cells. Sci. Rep. 8, 17559 (2018).

27. Wu, J. et al. Bacteriophage defends murine gut from Escherichia coli invasion via mucosal adherence. Nat. Commun. 15, 4764 (2024).

28. Bichet, M. C. et al. Bacteriophage uptake by mammalian cell layers represents a potential sink that may impact phage therapy. iScience 24, 102287 (2021).

29. Mund, A. et al. Deep Visual Proteomics defines single-cell identity and heterogeneity. Nat. Biotechnol. 40, 1231–1240 (2022).

30. Smith, K. et al. CIDRE: an illumination-correction method for optical microscopy. Nat. Methods 12, 404–406 (2015).

31. Hollandi, R. et al. nucleAIzer: A Parameter-free Deep Learning Framework for Nucleus Segmentation Using Image Style Transfer. Cell Syst. 10, 453–458.e6 (2020).

