## Supplementary figures for "Prevalent Gut Phages Encode Modular Adhesins Mediating Epithelial Binding and Endoplasmic Reticulum Trafficking"

**^2^ Doctoral School of Biology, University of Szeged, Szeged, Hungary**

**^3^ Department of Genetics, ELTE Eötvös Loránd University, Budapest, Hungary**

**^4^ Biological Barriers Research Group, Institute of Biophysics, HUN-REN Biological Research Centre, Szeged, Hungary**

**^5^ Single-Cell Technologies Ltd., Szeged, Hungary**

**^6^ Institute of AI for Health, Helmholtz Zentrum München, Neuherberg, Germany**

**^7^ Institute of Medical Microbiology and Immunology, University of Pécs, Pécs, Hungary**

**^8^ APC Microbiome Ireland & School of Microbiology, University College Cork, Co. Cork**

**^9^ Department of Medicine, University College Cork, Co. Cork**

**† These authors contributed equally.**

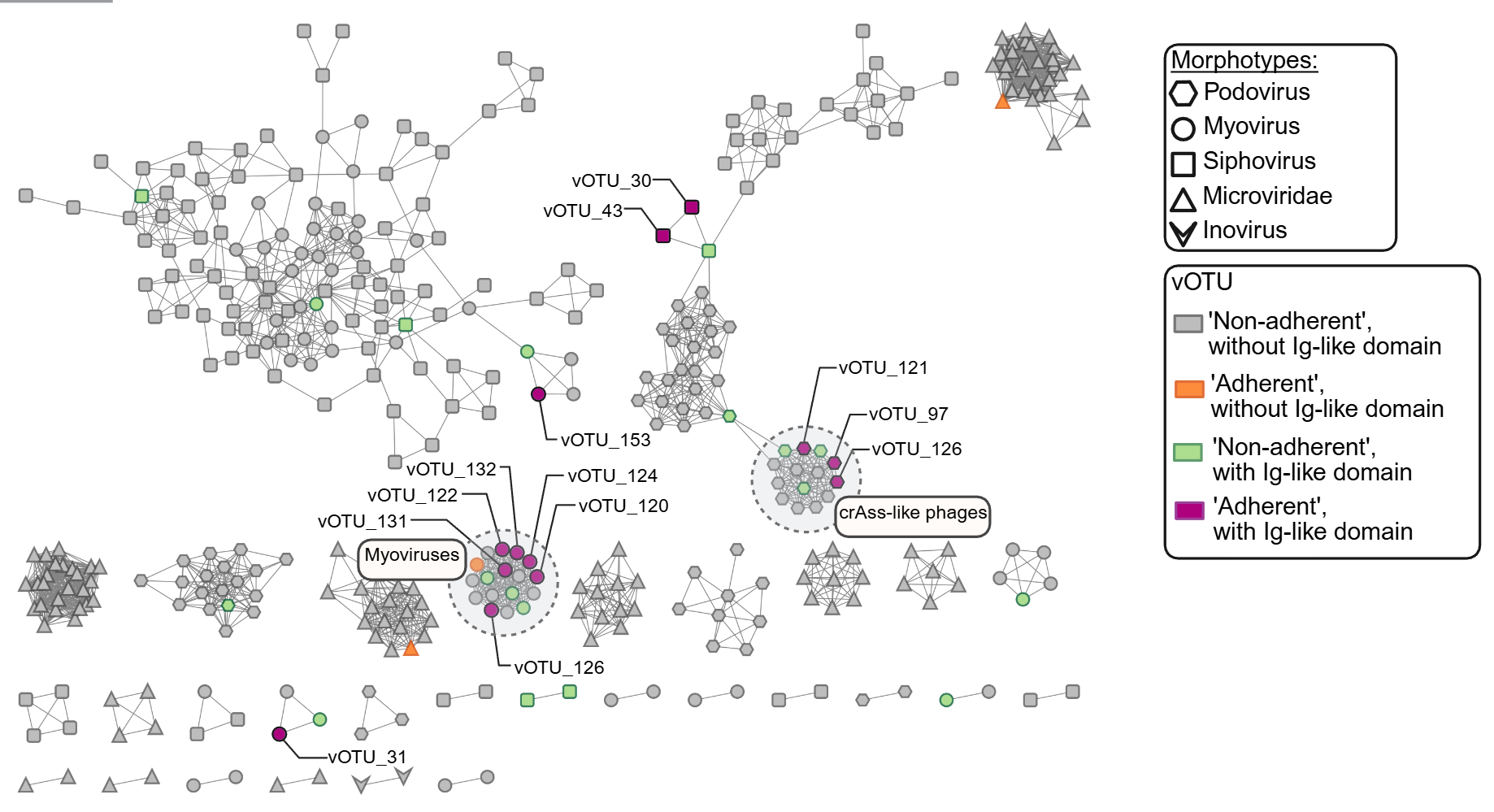

**Figure S1.** Protein-orthogroup–based network of 451 complete vOTUs identified with short-read sequencing of the fecal filtrate, the residual fraction, and the effluent fraction in the high-throughput selection experiment (see Methods). Node shapes represent phage morphotypes, and nodes are colour-coded by adherence phenotype and presence or absence of Ig-like domains. Singleton nodes are not shown. Experimental details are available in the Methods section, sequence data and taxonomy are in Tables S1 and S2, and the input for the network figure is in Table S7.

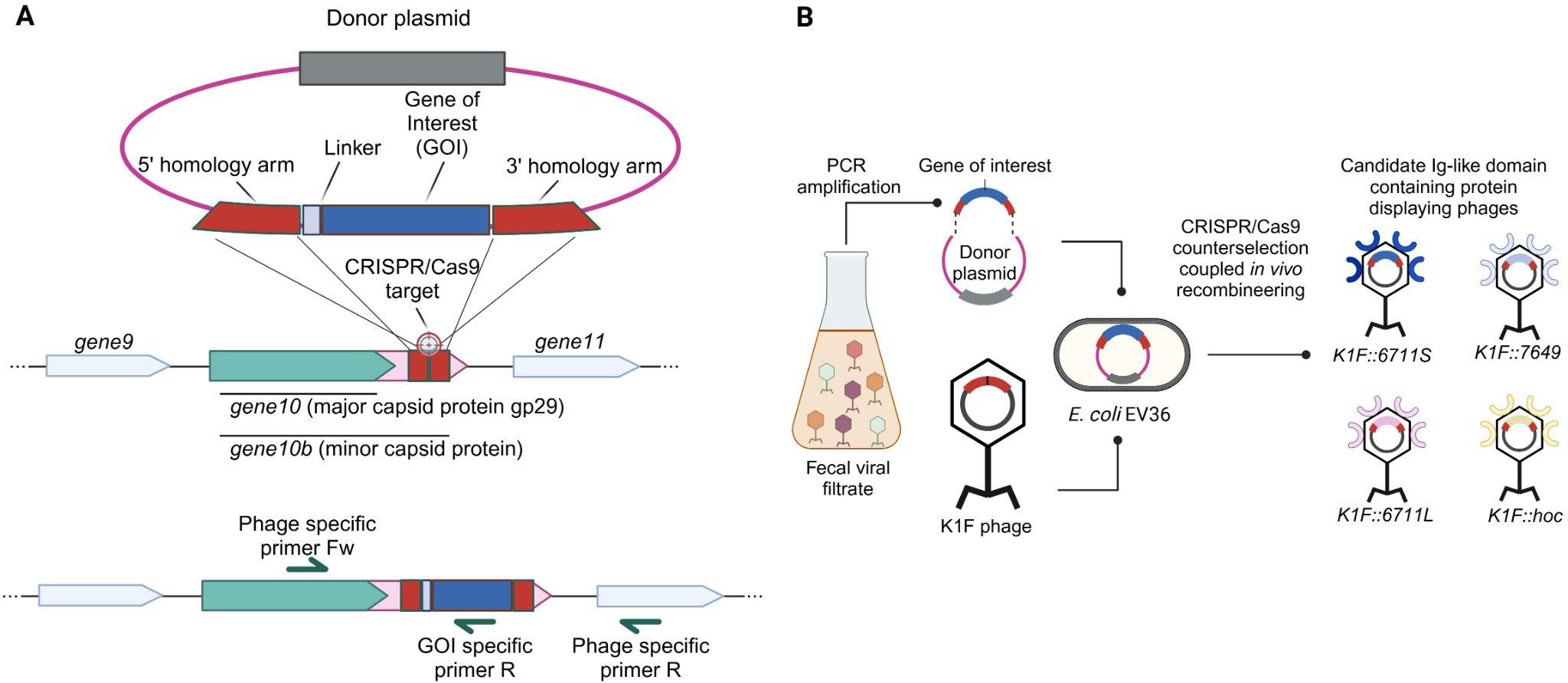

**Figure S2.** (A) Schematic representation of the recombination method adapted from Møller-Olsen and colleagues^38^ with modifications, used for the integration of the identified Ig-like domain-containing proteins and the Hoc protein (indicated as ‘Gene of Interest’) into the K1F phage genome. During the translational process, a -1 frameshift in *gene10* can produce an extended protein product, gene10b, which serves as the minor capsid protein displaying the fusion protein. (B) Schematic representation of the experimental procedure used to construct the Ig-like domain-displaying K1F phages. The genes of interest were PCR amplified, introduced by Golden Gate assembly into the donor plasmid, which was subsequently transformed into the host EV36 *E. coli* cells. The recombination step took place during the propagation of the K1F phage. After a CRISPR/Cas9-driven counterselection, the phages containing the gene of interest were identified by PCR using specific primers (Table S23).

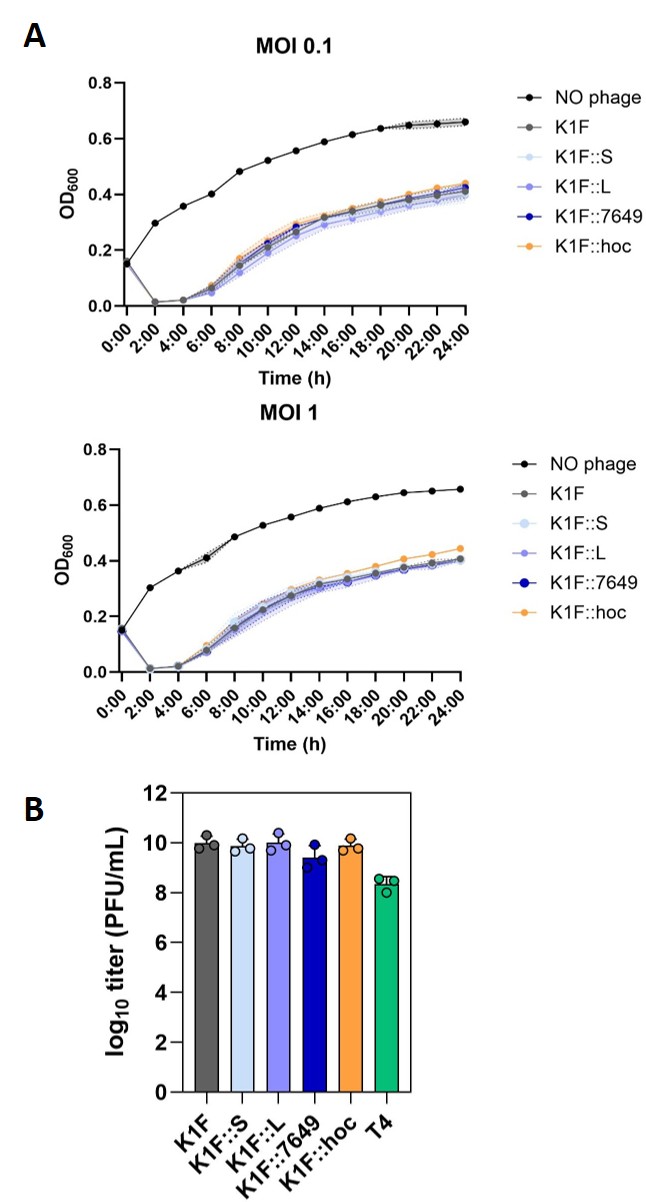

**Figure S3.** **Characterization of the engineered K1F phages. (A)** Growth curves (OD600) of *E. coli* EV36 strain in the absence and presence of wild-type or engineered K1F phages at MOI 0.1 and MOI 1, respectively. There is no significant difference in the growth of the EV36 strain in the presence of the engineered K1F variants compared to the wild-type K1F phage (see Table S11 for raw data). Mean and standard deviation of 3 replicates are shown. MOI - Multiplicity of infection, representing the ratio of phages to bacteria. **(B)** Comparing the titer of the engineered and wild-type K1F phages. Phage titer is shown as log_10_ PFU/mL. Mean and standard deviation are shown (n = 3).

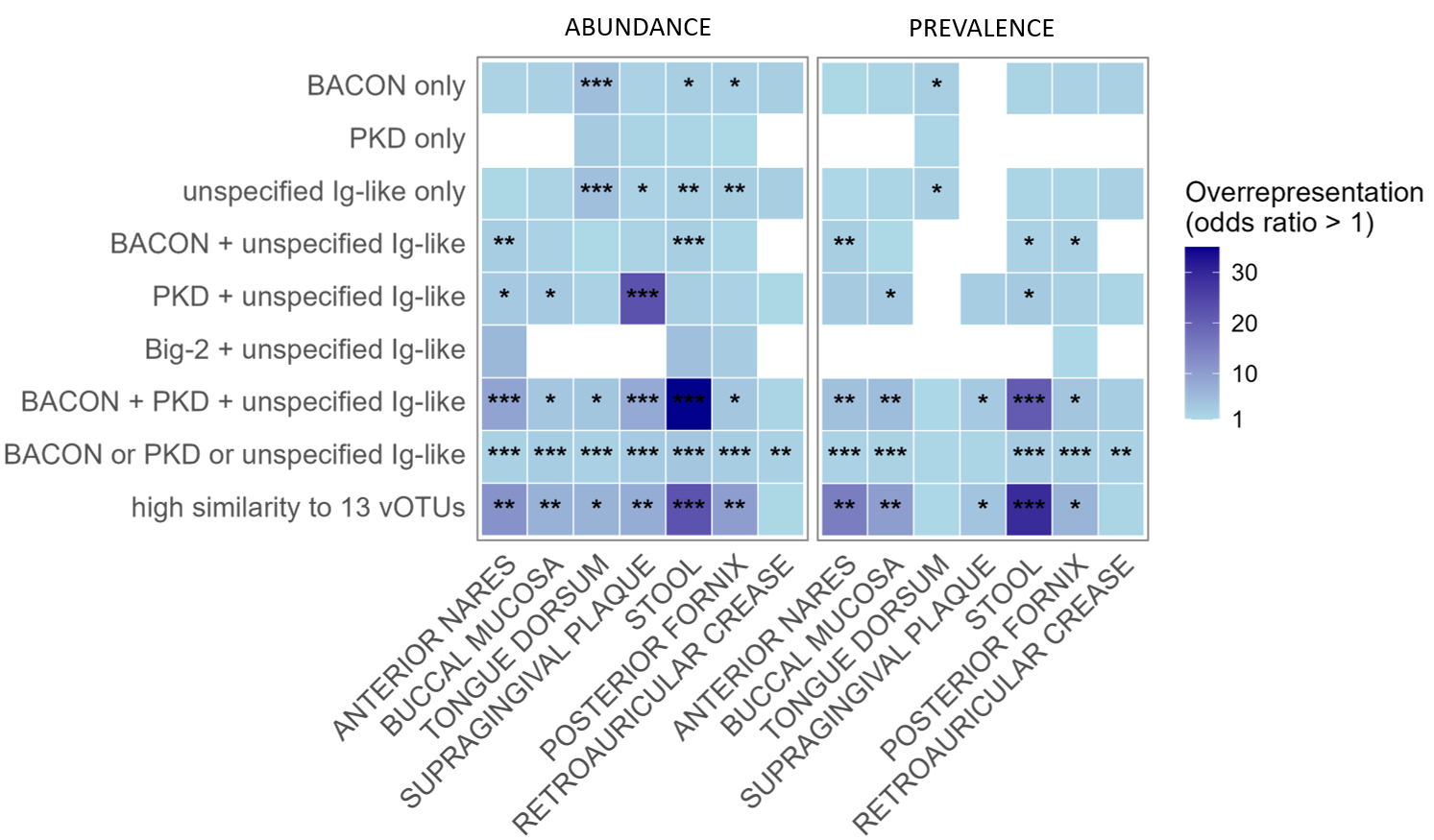

**Figure S4. Phages harbouring BACON, PKD, or unspecified Ig-like domains, identified in the adherent vOTUs are overrepresented among highly abundant and highly prevalent Bacteroidota-infecting phages across different mucosal body sites.** Blue intensity indicates the degree of overrepresentation, based on odds ratios calculated using Fisher’s exact test. Statistically significant enrichments are marked with asterisks (*p < 0.05, **p < 0.01, ***p < 0.001), with p-values adjusted using the Benjamini–Hochberg correction.

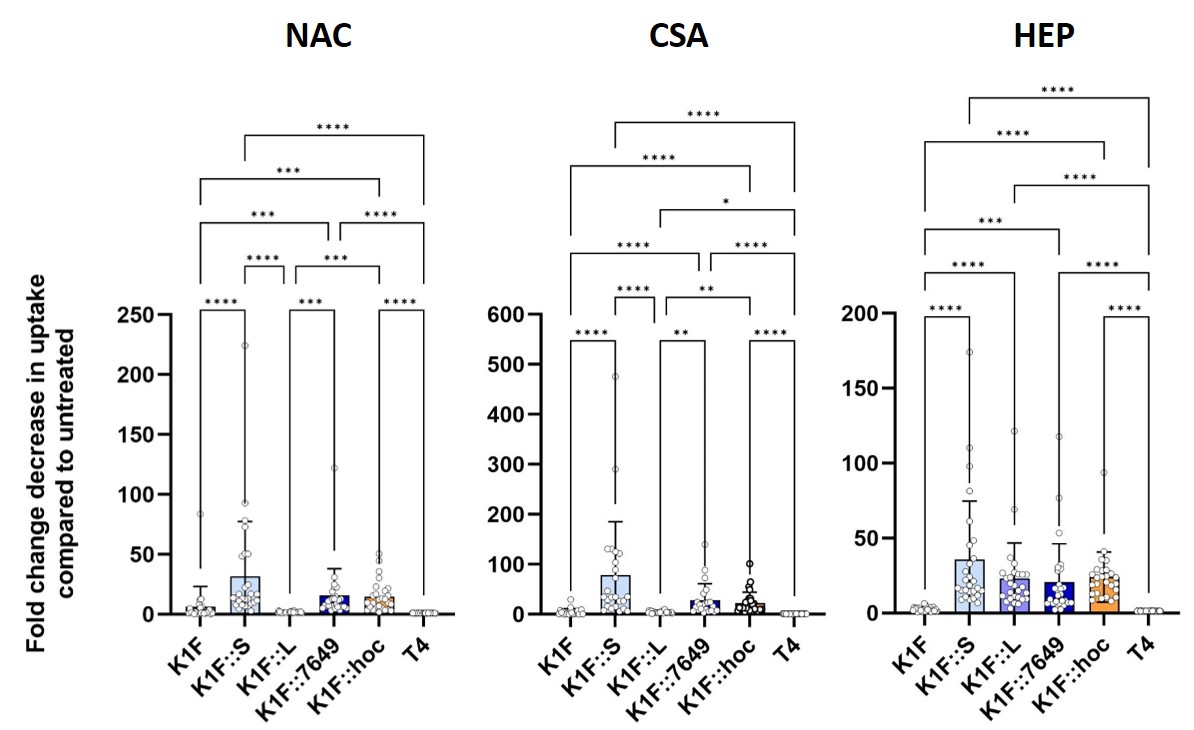

**Figure S5.** **Fold change decreases in uptake of the wild-type and engineered K1F phages, and the T4 phage by HT29-MTX epithelial cells in response to different treatments of the epithelial cell surface.** Treatments: NAC - N-acetylcysteine, CSA - cyclosporin A, HEP - heparinase I. Fold change was determined from area under curve (AUC) values, which were calculated from the percentage of cells with intracellular phages over time. Mean and standard deviation are shown. Each dot represents one field of view. Stars indicate p-values calculated from Kruskal-Wallis test. For details see Methods and Table S21.

.

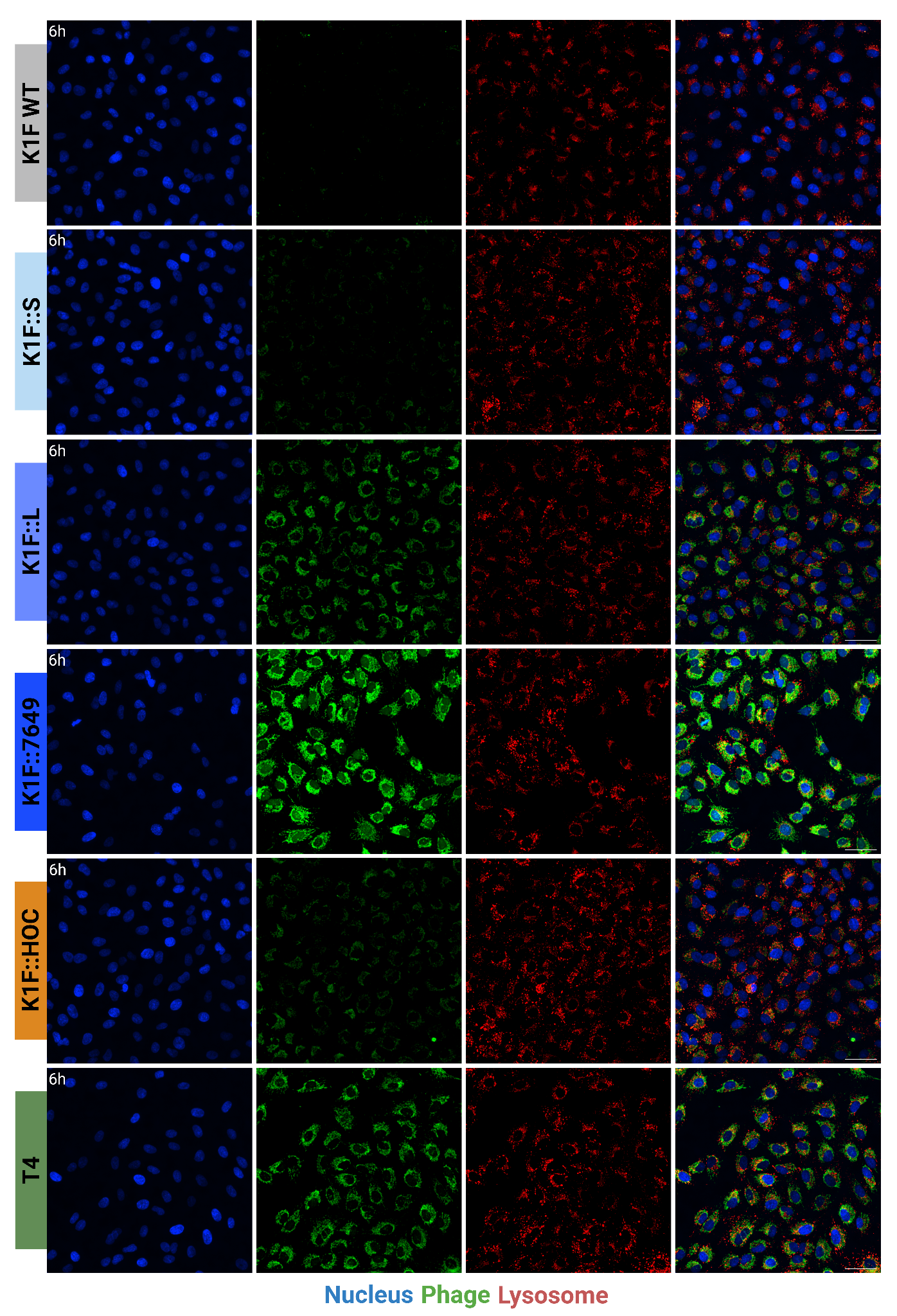

**Figure S6. High content confocal microscopy images showing the colocalization of the wild-type and engineered K1F phages with lysosomes in A549 lung epithelial cells after 6 hours.** Cell nucleus was stained with Hoechst dye (blue), phages were labeled with SYBR™ Gold dye (green), while lysosomes were labeled with Lysotracker^TM^ Deep Red dye (for details, see Methods). Scale bar = 50 µm.

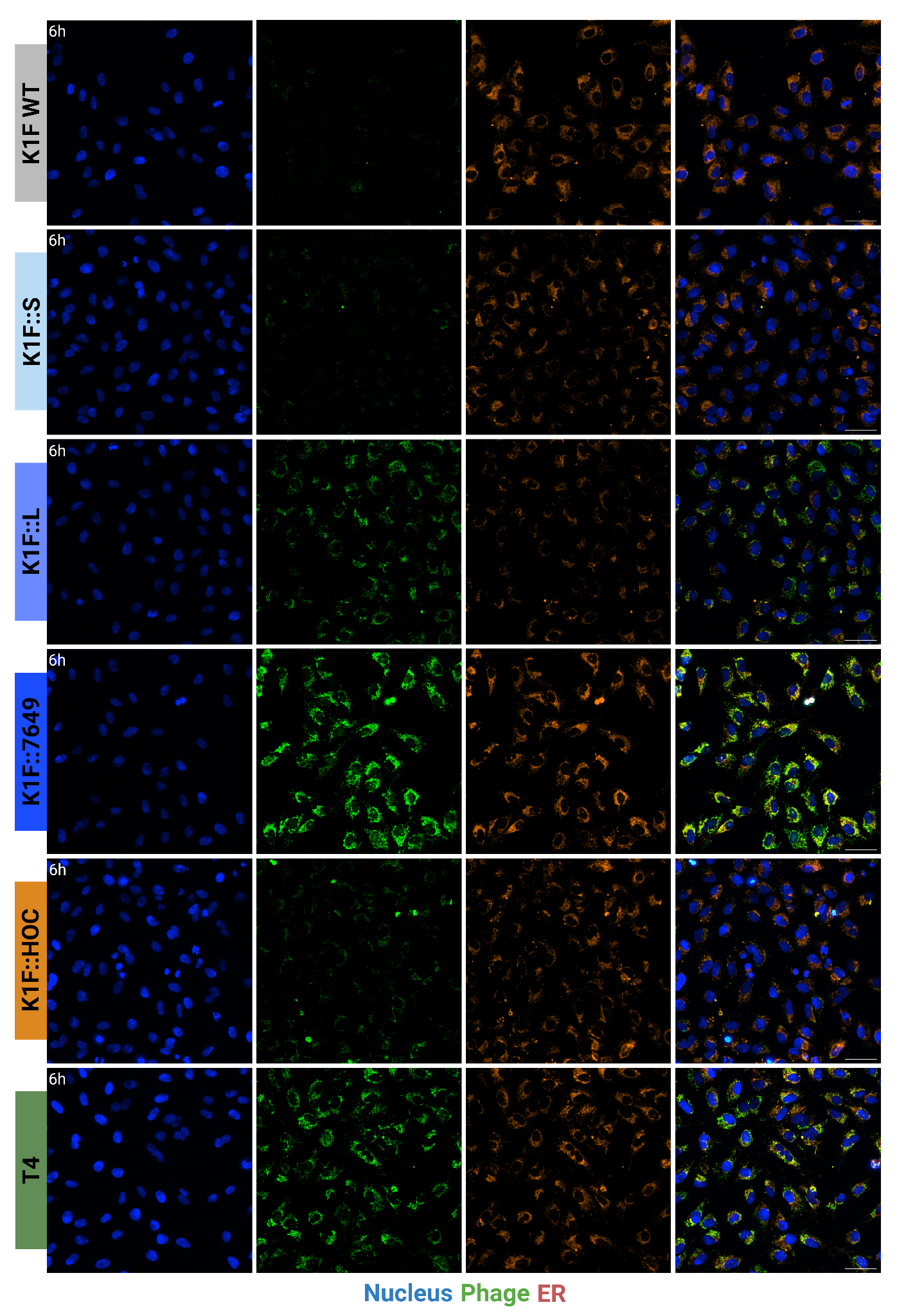

**Figure S7. High content confocal microscopy images showing the colocalization of the wild-type and engineered K1F phages with endoplasmatic reticulums in A549 lung epithelial cells after 6 hours.** Cell nucleus was stained with Hoechst dye (blue), phages were labeled with SYBR™ Gold dye (green), while endoplasmatic reticulums were labeled with ER-Tracker^TM^ Red dye (for details, see Methods). Scale bar = 50 µm.

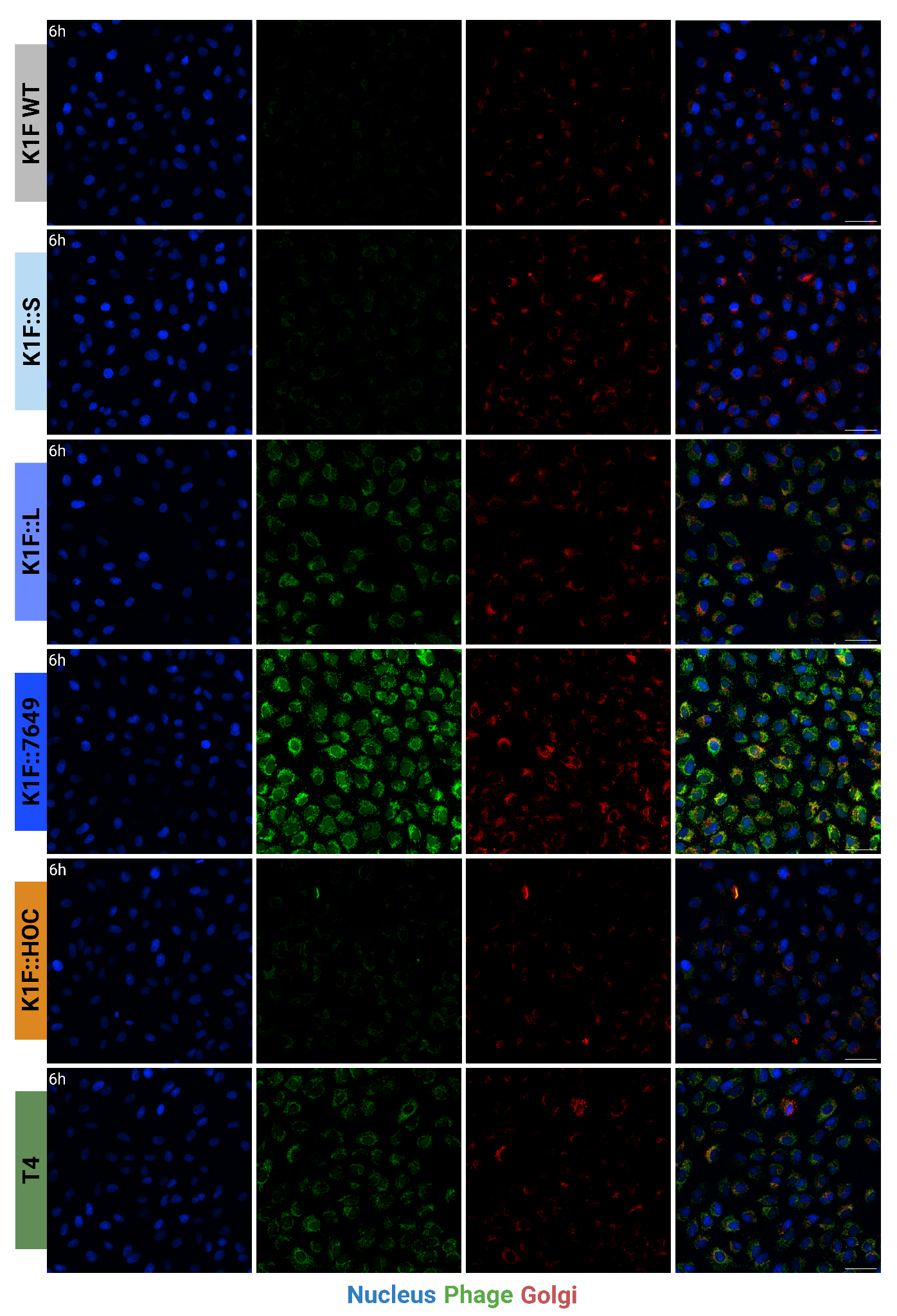

**Figure S8. High content confocal microscopy images showing the colocalization of the wild-type and engineered K1F phages with golgi apparatus in A549 lung epithelial cells after 6 hours.** Cell nucleus was stained with Hoechst dye (blue), phages were labeled with SYBR™ Gold dye (green), while golgi apparatus were labeled with BODIPY^TM^ TR Ceramide dye (for details, see Methods). Scale bar = 50 µm.
